# Neurotensin-Neurotensin Receptor 2 signaling in adipocytes regulates food intake through ceramide metabolism

**DOI:** 10.1101/2024.02.07.579397

**Authors:** Wei Fu, Yue Yang, Xiao Guo, Qifan Gong, Xiaofeng Zhou, Liying Zhou, Cenxi Liu, Zhi Zhang, Jisun So, Yufeng Zhang, Lin Huang, Guangxing Lu, Chuanyou Yi, Qichu Wang, Chenyu Fan, Chao Liu, Jiaxing Wang, Haiyi Yu, Yimin Zhao, Tao Huang, Hyun Cheol Roh, Tiemin Liu, Huiru Tang, Jianping Qi, Ming Xu, Yan Zheng, He Huang, Jin Li

**Affiliations:** State Key Laboratory of Genetic Engineering, School of Life Sciences, Institute of Metabolism and Integrative Biology, Human Phenome Institute and Zhongshan Hospital, Fudan University, Shanghai, 200438, China; Department of Endocrinology, The First Affiliated Hospital and Clinical Medicine College, Henan University of Science and Technology, Luoyang 471003, China; National Center for Clinical Research of Metabolic Diseases, Luoyang Center for Endocrinology and Metabolism, Luoyang 471003, China; Diabetic Nephropathy Academician Workstation of Henan Province, Luoyang 471003, China; Department of Biochemistry and Molecular Biology, Indiana University School of Medicine, Indianapolis, IN 46202, USA; Department of Cardiology and Institute of Vascular Medicine, Peking University Third Hospital; State Key Laboratory of Vascular Homeostasis and Remodeling, Peking University; NHC Key Laboratory of Cardiovascular Molecular Biology and Regulatory Peptides, Peking University, Beijing, China; Department of Epidemiology and Biostatistics, School of Public Health, Peking University, Beijing, China

**Keywords:** Neurotensin, Adipocyte, Non-shivering thermogenesis, Food intake, Ceramide

## Abstract

Neurotensin (NTS) is a secretory peptide produced by the lymphatic endothelial cells (LEC). Our previous study revealed that NTS suppressed the activity of brown adipose tissue via the interactions with NTSR2. In the current study, we found that the depletion of *Ntsr2* in the white adipocytes upregulated food intake, while the local treatment of NTS suppressed the food intake. Mechanistic study revealed that the suppression of NTS-NTSR2 signaling enhanced the phosphorylation of ceramide synthetase 2 (CerS2), increased the abundance of its products ceramide C20-C24 and downregulated the production of GDF15 in the white adipose tissues, which was responsible for the elevation of food intake. With four populations of different age and ethnic background, we discovered a potential causal and positive correlation between ceramide C20-24 and food intake in human. Our study identified that NTS-NTSR2 signaling can perform the neurological regulation via controlling the production of ceramide in the white adipocytes.

## INTRODUCTION

Adipose tissue is one of the largest organs and a major regulator of metabolic homeostasis in mammals. Though its primary function is lipid storage, it affects metabolism via various routes. Several studies have demonstrated that the adipose tissue is an important endocrine organ. Adipose tissue derived secretary proteins, from either the adipocytes or stromal vascular fraction (SVF), tightly modulate the metabolic activity of many organs. One such well known factor is leptin, which is an adipose tissue derived secretory protein that regulates multiple aspects of metabolism via interactions with the leptin receptor (*1, 2*). Adipose tissue also can affect the physiological function of skeletal muscle via the secretory protein myostatin (*3*). In addition to endocrine activity, adipose tissue derived secretory proteins can regulate metabolic homeostasis in an autocrine or paracrine signaling as well (*4*).

Neurotensin (NTS) is a 13 amino acid secretory peptide with diverse functions that act upon both the central nervous system (CNS) and peripheral tissues. Due to its very short half-life in the circulation (*5, 6*), the native NTS can only exert its functions via the paracrine or autocrine mode. NTS was previously known to be produced by neurons and intestinal N cells (*7–9*). Two G protein coupled receptors (GPCR), NTSR1 and NTSR2, have been discovered to specifically interact with NTS (*10, 11*). The type I membrane glycoprotein Sortilin (SORT1 or NTSR3) also interacts with NTS, though its downstream signaling is largely uncharacterized (*12–14*). Many of these studies attributed the decreases in food intake to the local and direct interaction between NTS and cells in the CNS, as intracerebroventricular (ICV) injection of NTS was sufficient to reduce food intake (*15–18*). The feeding behavior changes induced by NTS treatment is likely mediated by NTSR1, though the role of NTSR2 cannot be excluded (*19–22*).

As far as we know, it is unknown whether NTS can regulate food intake through direct interactions with peripheral tissues or organs in the physiological conditions. One possible reason is that for a long period of time, people believe that NTS is only produced by either CNS or intestinal N cells. Though the results of ICV injection indicated that NTS could regulate food intake via direct interactions with CNS, it did not rule out the possibility that NTS could also regulate food intake via interacting with peripheral tissue in physiological conditions.

With single cell RNA-seq analysis, we have determined that NTS was also produced specifically by the lymphatic endothelial cells (LECs) as an anti-thermogenic peptide in the brown adipose tissue (*23*) and a pro-lipid absorption peptide regulating the development of atherosclerosis in the intestine (*24–26*). *Ntsr2*, but not *Ntsr1*, is widely expressed in brown adipocytes, beige adipocytes and white adipocytes (*27*). Transient knock-down experiments have indicated that NTS exerts its anti-thermogenic effects via the interactions with NTSR2 in brown adipose tissue (BAT) through paracrine signaling. However, the cell type specific effects and long-term physiological functions of NTSR2, especially how it functions in the white adipocytes, are still unknown.

Ceramide is a kind of lipids with diverse biological functions. The metabolism of ceramide is extremely complicated, as these lipids can be synthesized and degraded by a large collection of enzymes in various pathways (*28*). It has been reported that the depletion of ceramide synthetase 2 (*CerS2*) led to the downregulation of its products ceramide C20-C24 and upregulation of C16 due to the compensatory effects in the liver (*29–33*). In addition, it has been demonstrated that ceramide C20-C24 is a negative regulator of the unfolded protein response (UPR) (*34, 35*). The UPR, as a part of the integrative stress response, may affect the production of a secretory protein Growth differentiation factor 15 (GDF15) (*36, 37*). On one hand, GDF15 treatment suppresses food intake and decreases body weight via interactions with its receptor, GFRAL, in the area postrema/nucleus of the solitary tract (AP/NTS) region of the CNS; On the other hand, depletion of *Gdf15* or *Gfral* gene leads to increases in food intake and body weight (*38–47*). However, the mechanism to regulate the production and physiological functions of ceramide C20-C24 has not been elucidated.

In the current study, we established an brown/beige adipocyte specific KO mouse model for *Ntsr2* (*Ucp1-cre*::*Ntsr2*^flox/flox^, referred to *Ntsr2* BKO mice), which exhibited an elevated energy expenditure and lower body weight when fed by a high fat diet (HFD). In contrast, the adipocyte specific *Ntsr2* depletion (*Adipoq-cre*::*Ntsr2*^flox/flox^, referred to *Ntsr2* AKO mice) presented the elevation of food intake, but the local treatment of NTS induced remarkable decreases of food intake in a NTSR2 dependent manner. The NTS-NTSR2 signaling controlled the synthesis of ceramide C20-C24, but not C16, by suppressing the phosphorylation of CerS2 via RhoA but not canonical CK2 kinase in the white adipocytes. The suppression of ceramide C20-C24 production in the adipose tissue by depleting *CerS2* gene downregulated food intake, but the local treatment of ceramide C20-C24 *in vivo* upregulated food intake. The ceramide-UPR dependent production of GDF15 mediated the cross-talk between the NTS-NTSR2 signaling in the adipose tissue and food intake. With two newly established populations, we confirmed that the serum abundance of ceramide C20-C24 was positively correlated to the energy intake in both children and adults. With mendelian randomization analysis, we discovered the potential causal relation between ceramide C20-C24 and food intake in human. The current study provides the first evidence for NTS-NTSR2 signaling to control the food intake by directly regulating the ceramide metabolism in adipose tissues.

## RESULTS

### Brown/beige Adipocyte specific depletion of *Ntsr2* increases the energy expenditure

A conditional knockout mouse model for *Ntsr2* was established by knocking in the *LoxP* cassette before the first exon and after the last exon of *Ntsr2* gene. By crossing the *Ntsr2*^flox/flox^ mice with a *Ucp1-cre* mouse, the *Ntsr2* BKO mouse model was established for phenotypic characterization (**Supplementary Figure 1A**). The *Ntsr2* BKO efficiency and specificity in the brown adipose tissue was validated by RT-qPCR (**Supplementary Figure 1B**). The body weight and food intake of *Ntsr2* BKO mice and control mice (*Ntsr2*^flox/flox^ mice) fed by a chow diet was comparable (**Supplementary Figure 1C-D**). Notably, *Ntsr2* BKO mice have a lower body weight than control mice when they were fed by HFD (**Supplementary Figure 1E**). More lean mass and less fat mass were observed in the *Ntsr2* BKO mice (**Supplementary Figure 1F**). The *Ntsr2* BKO mice also had a better performance in the glucose tolerance test (GTT) (**Supplementary Figure 1G**) and insulin tolerance test (ITT) (**Supplementary Figure 1H**).

The Comprehensive Lab Animal Monitoring System (CLAMS) analysis revealed *Ntsr2* BKO mice presented a higher O2 consumption rate (**Supplementary Figure 1I**) and CO2 production rate (**Supplementary Figure 1J**), indicating the enhancement of energy expenditure. *Ntsr2* BKO mice also had increased expression of thermogenic genes in the brown adipose tissue (BAT) but not inguinal white adipose tissue (iWAT) (**Supplementary Figure 1K-L**). In contrast, the physical activity and fecal energy between *Ntsr2* BKO mice and control mice were comparable (**Supplementary Figure 1M-N**). It is widely observed that the elevated thermogenesis only had effects on the body weight of obese but not young and lean mice (*48, 49*). These results indicated that NTSR2 in the BAT has significant impacts on thermogenesis and development of diet induced obesity (DIO).

### NTS-NTSR2 signaling in the white adipocytes regulated food intake

By employing the transcriptomic analysis of adipocyte-specific mRNA isolated by translating ribosome affinity purification (TRAP) (Roh et al., 2020; Roh et al., 2017; Roh et al., 2018), we found that the *Ntsr2* expression was dramatically upregulated by HFD specifically in the epididymal WAT (eWAT) (**Figure 1A**). We therefore further explored the functions of NTSR2 in white adipocytes. By crossing the *Ntsr2*^flox/flox^ mice with an *Adipoq-cre* mouse, the *Ntsr2* AKO mouse model was established for phenotypic characterization (**Figure 1B**). The *Ntsr2* depletion efficiency and specificity in the adipose tissues and adipocytes was validated by RT-qPCR (**Figure 1C, Supplementary Figure 2A-B**). The body weight of *Ntsr2* AKO mice and control mice (*Ntsr2*^flox/flox^ mice) fed by a chow diet was comparable (**Supplementary Figure 2C**). But the *Ntsr2* AKO mice have a higher body weight than control mice when fed by HFD (**Figure 1D**). There is a trend of increased fat mass and lean mass in the *Ntsr2* AKO mice (**Supplementary Figure 2D**). The *Ntsr2* AKO mice also had a worse performance in the GTT (**Figure 1E**) and ITT (**Figure 1F**).

**Figure 1.**
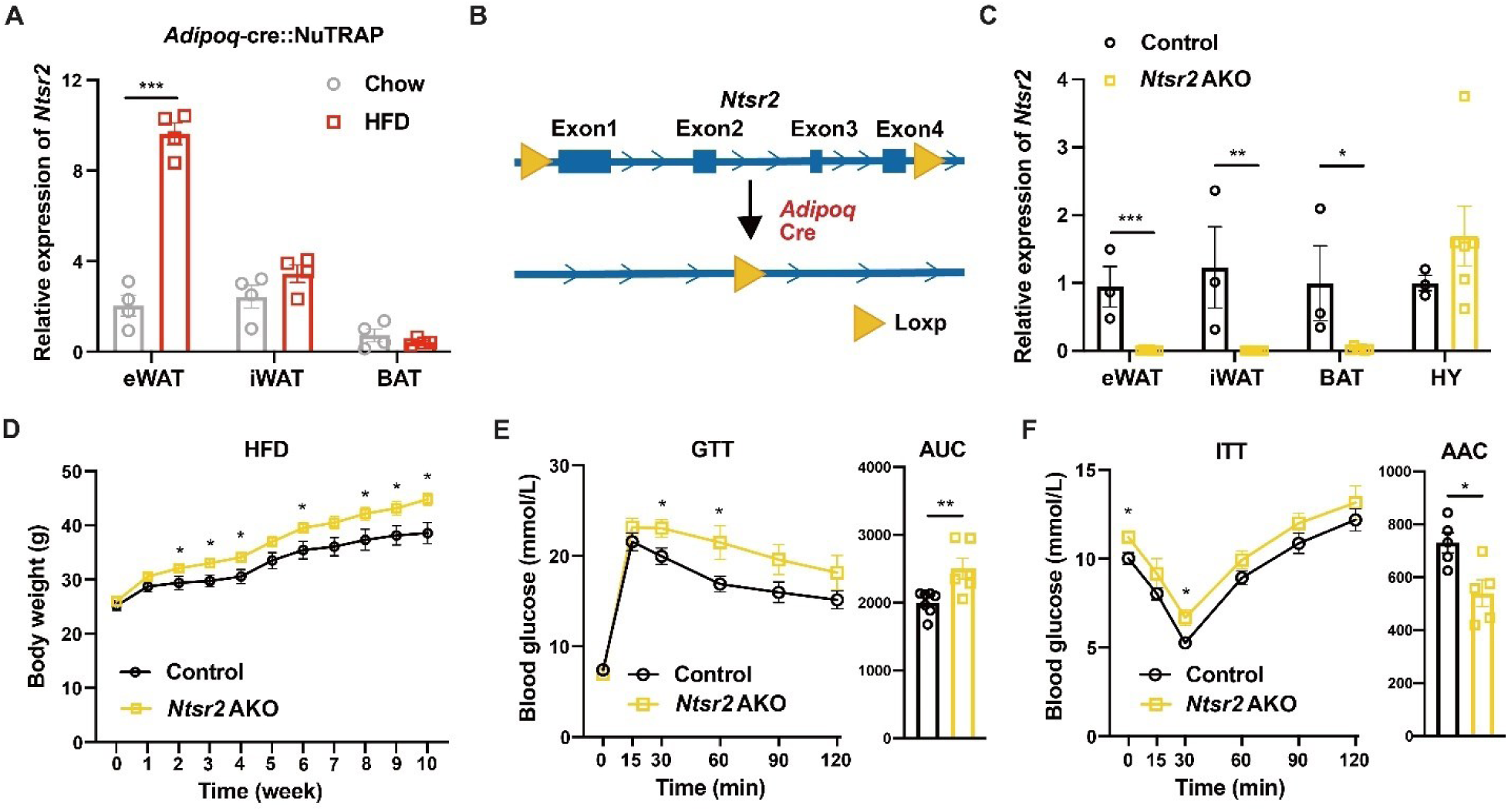
The depletion of *Ntsr2* in the adipocytes induced the elevation of body weight. **A.** The expression of *Ntsr2* in the adipocyte-specific transcriptome of lean and obese mice; N=4; **B.** The establishment of adipocyte specific *Ntsr2* KO mouse model; **C.** The KO efficiency and specificity of the *Ntsr2* gene; N=3-6; **D-F.** The Body weight (**D**), GTT (**E**) and ITT (**F**) of mice fed by HFD; N=5-7. *, p<0.05; **, p<0.01; ***, p<0.001.

We then explored the reasons for the body weight changes in the *Ntsr2* AKO mice. It is known that the impaired lipolysis might be an important biological process contributing to the development of DIO. However, we found that *Ntsr2* AKO rather led to the elevation of lipolysis related proteins p-HSL and ATGL in the eWAT (**Supplementary Figure 2E**). The concentration of non-esterified fatty acids (NEFA) in serum was also elevated in *Ntsr2* AKO mice upon the challenge of fasting (**Supplementary Figure 2F**). To investigate whether the changes of lipolysis were due to the cell autonomous effects of NTS-NTSR2 signaling, we treated the primary adipocytes derived from the SVF with recombinant NTS peptides. NTS treatment decreased the abundance of lipolysis related protein ATGL and p-HSL (**Supplementary Figure 2G**). It also prevented lipolysis, determined by the reduced release of NEFA from adipocytes (**Supplementary Figure 2H**). Notably, the changes of p-HSL and ATGL protein abundance and NEFA release, upon treatment with NTS, were absent in adipocytes from *Ntsr2* AKO mice (**Supplementary Figure 2I-J**). Thus, we concluded that the changes of lipolysis are not responsible for the upregulation of body weight in the *Ntsr2* AKO mice.

We then explored the status of energy metabolism in these mice. Similar to *Ntsr2* BKO mice, the *Ntsr2* AKO mice also presented a trend of increased thermogenesis (**Supplementary Figure 3A-B**). But in contrast to the *Ntsr2* BKO mice, the food intake of *Ntsr2* AKO mice was remarkably increased (**Figure 2A and Supplementary Figure 3C**). Notably, the *Ntsr2* AKO mice on pair feeding of HFD showed lower body weight as well as fat mass weight (**Figure 2B**). The pair-fed *Ntsr2* AKO mice also had a better performance in the GTT (**Figure 2C**) and ITT (**Figure 2D**).

**Figure 2.**
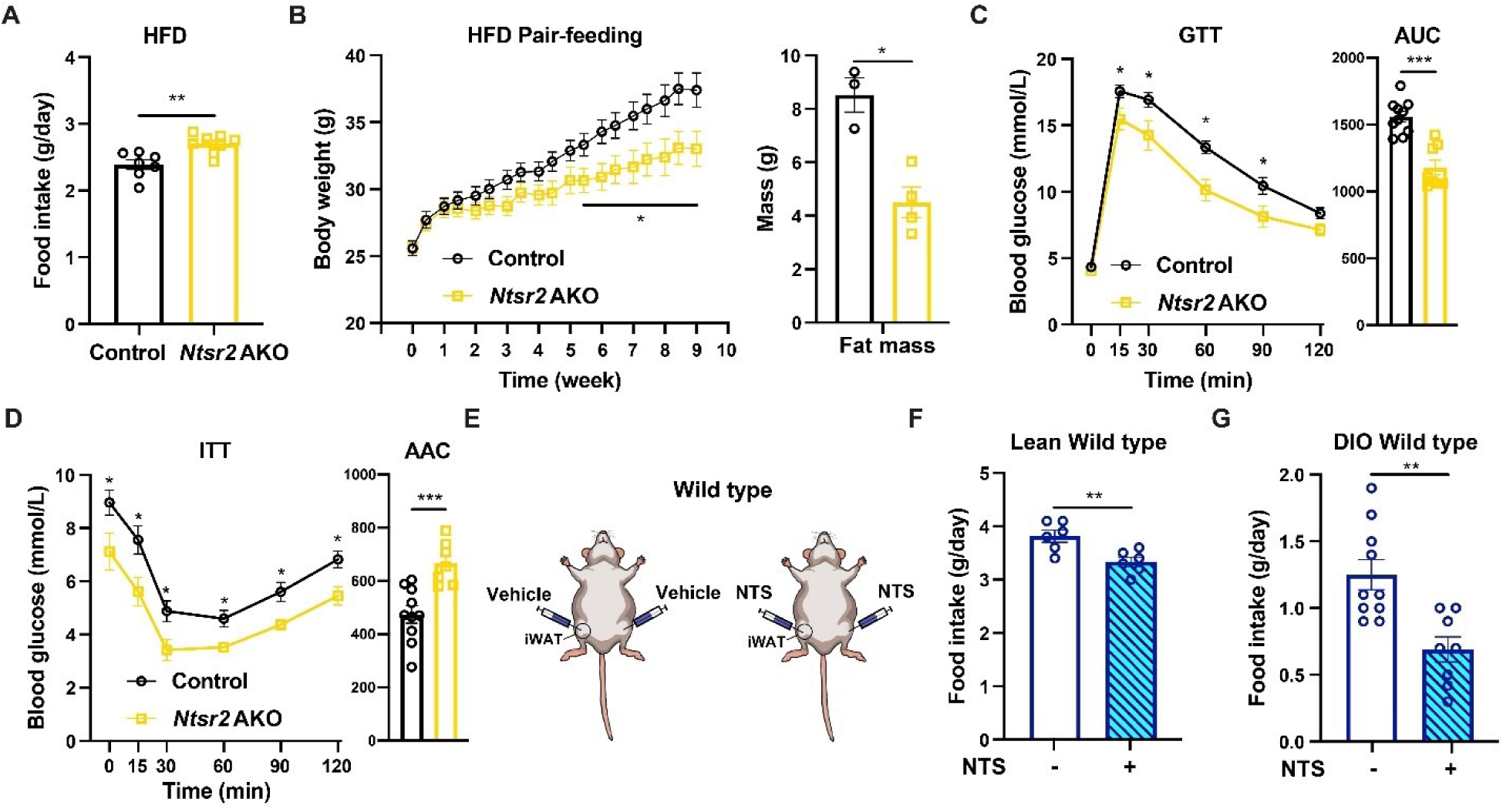
NTS-NTSR2 signaling in the adipose tissue regulated food intake. **A.** The food intake of control and *Ntsr2* AKO mice; N=7-8; **B-D.** The changes of body weight (**B**), GTT (**C**) and ITT (**D**) of mice pair-fed by HFD. N=6-10; **E.** The illustration of local NTS treatment; **F-G.** The food intake of wild-type lean (**F**) or obese (**G**) mice upon NTS treatment. N=6-12. *, p<0.05; **, p<0.01; ***, p<0.001.

Based on the data related to the changes of food intake in the *Ntsr2* AKO mice, we tested the local effects of recombinant NTS on the iWAT *in vivo* with the hydrogel containing recombinant NTS peptide (**Supplementary Figure 3D**). The hydrogel was injected to the iWAT of wild type C57 BL6/J mice (**Figure 2E**). Comparing to the treatment of the hydrogel itself, the local release of NTS in the WAT led to the decreases of food intake in both lean (**Figure 2F**) and obese mice (**Figure 2G**). Especially, the local release of NTS has no effects on the food intake of *Ntsr2* AKO mice (**Figure 3E**), demonstrating that NTS regulated the food intake via the interactions with NTSR2 in the adipocytes. These results suggested that the NTS-NTSR2 signaling regulated the food intake via the direct effects on the adipose tissue.

**Figure 3.**
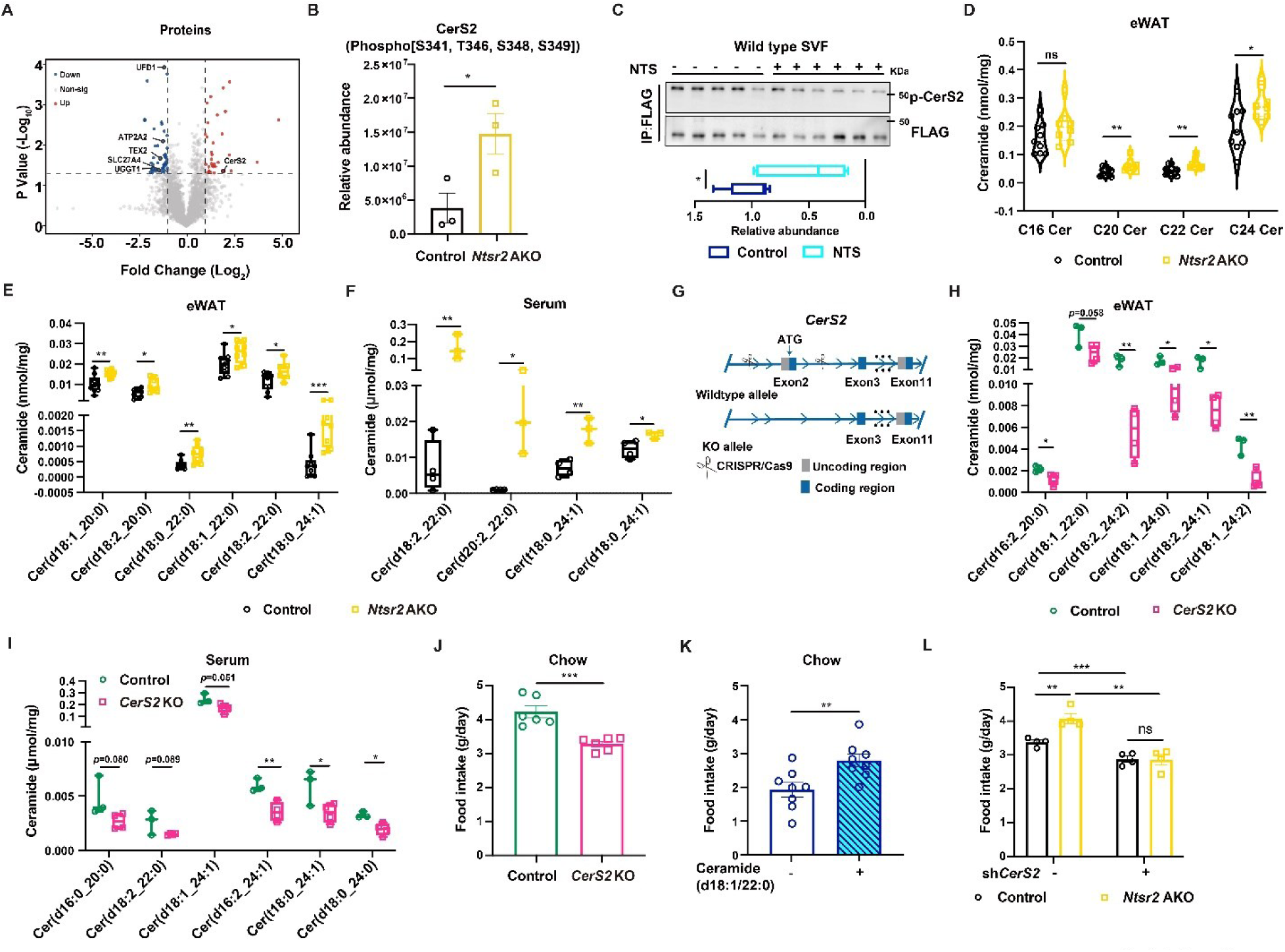
NTS-NTSR2 signaling regulated the ceramide metabolism in the WAT. **A.** A volcano plot of phospho-proteomics data; **B.** The relative abundance of phosphorylated CerS2 (p-CerS2). N=3; **C.** The level of p-CerS2 detected by pull-down and western blot in the primary adipocytes. N=5-6; **D-E.** The concentration of ceramide C16-C24 (**D**) or individual ceramide (**E**) in the WAT of control or *Ntsr2* AKO mice. N=6; **F.** The concentration of individual ceramide in the serum of control or *Ntsr2* AKO mice. N=6; **G.** The illustration of *CerS2* KO mice; **H-I.** The concentration of individual ceramide in the WAT (**H**) or serum (**I**) of control or *CerS2* KO mice. N=3; **J-K.** The food intake of control and *CerS2* KO mice (**J**) or vehicle versus ceramide C22 treated mice (**K**). N=6; **L.** The food intake of the control and *Ntsr2* AKO mice with the knocking-down of *CerS2*. N=4. *, p<0.05; **, p<0.01; ***, p<0.001.

### NTS-NTSR2 signaling affected the metabolism of ceramide in adipocytes

We then explored the mechanism for the regulation of food intake by the NTS-NTSR2 signaling in the adipose tissues. We have reported that NTS-NTSR2 signaling regulated the MEK-ERK in the BAT, but this phenomenon was absent in the eWAT of *Ntsr2* AKO mice (**Supplementary Figure 4A**). The phospho-proteomics analysis on the eWAT from lean control and *Ntsr2* AKO mice with comparable body weight identified the significant changes of phosphorylation for multiple proteins (**Figure 3A**). Specifically, *Ntsr2* depletion induced the elevation of phosphorylation in ceramide synthetase 2 (CerS2) protein (**Figure 3B**). Other ceramide synthetases were not detected in the phospho-proteomics analysis. To confirm the regulation of phosphorylated-CerS2 (p-CerS2) by NTS-NTSR2 signaling, we overexpressed the *CerS2* gene with the Flag-HA tag in the N terminal (**Supplementary Figure 4B-C**). By pulling down CerS2 with the FLAG tag and performing immunoblot with a pan phosphor-Ser/Thr antibody, we identified that NTS suppressed the phosphorylation of CerS2 in the primary adipocytes (**Figure 3C**). In contrast, the inhibitor of CK2 (*29*), which has been identified as a kinase for the phosphorylation of CerS2 in 293T cells, has no obvious effects on the phosphorylation of CerS2 (**Supplementary Figure 4D**). The different molecular weight of the bands for the native and phosphorylated CerS2 was potentially due to glycosylation, which was consistent with what has been reported (*29*).

With the untargeted lipidomics assay, we observed the increase of C20-C24 ceramides as the products of CERS2, but not other species of ceramide especially C16, in the eWAT of *Ntsr2* AKO mice (**Supplementary Figure 4E**). In contrast, we did not observe the changes of sphingosine or sphingomyelin (**Supplementary Figure 4F**). These results were further validated by the targeted lipidomics assay, which was specifically designed for the detection of ceramide with various length of acyl chains (*32*) (**Figure 3D; The technical details of targeted lipidomics were described in the “*METHODS*”**). The top-5 differentially detected ceramides were presented at **Figure 3E**. Although the total concentration of C20-C24 ceramides was not dramatically changed in the serum, many C20-C24 ceramides were observed among the top differentially detected ceramides (**Figure 3F, Supplementary Figure 4G**). In the phospho-proteomics analysis, we also discovered that the depletion of *Ntsr2* regulated the phosphorylation of ceramide transporter (CERT) (**Supplementary Figure 4H**). As it has been reported that CERT was correlated to the abundance of C16 ceramide (*50*), it was unlikely to directly contribute to the effects of NTS-NTSR2 signaling in the current context.

We then studied the physiological functions of ceramide C20-C24 *in vivo*. It has been reported that the homozygous depletion *CerS2* was detrimental to the overall health of mice (*31*). In the current study, the depletion of one copy of *CerS2* gene (**Figure 3G and Supplementary Figure 5A**) led to the significant decreases of ceramide C24, trends of decreases of C20-C22, but without dramatic changes of C16 in both the eWAT and serum (**Supplementary Figure 5B-C**). The top-5 differentially detected ceramides in the WAT and serum were presented at **Figure 3H-I**. The expression of *CerS5* and *CerS6* was also unchanged in the adipose tissue with the haploid depletion of *CerS2* (**Supplementary Figure 5D**). In contrast, the liver of mice presented the significant decreases of individual ceramide C24, increases of individual C16 and upregulation for the expression of *CerS5* and *CerS6* (**Supplementary Figure 5E-F**), which was consistent to the literature (*31, 32*). The changes of C20-24, but not C16, in the WAT can be attributed to the differences of transcriptional regulation for *CerS5* and *CerS6* between liver and WAT.

The *Cers2* KO mice showed a decrease of food intake (**Figure 3J**). In contrast, the local treatment of ceramide C22, but not C16, in the WAT upregulated the food intake of mice *in vivo* (**Figure 3K and Supplementary Figure 5G**). Especially, the knocking-down of *CerS2* via injecting adeno associated virus (AAV) containing shRNA to the iWAT decreased the abundance of ceramide C20-24 without effects on C16 locally (**Supplementary Figure 5H-I**). The knocking-down of *CerS2* also normalized the upregulation of food intake induced by *Ntsr2* AKO (**Figure 3L**). These data indicated that the ceramide metabolism in the adipocyte contributed to the NTS-NTSR2 signaling regulated food intake.

### NTS-NTSR2 signaling affects the level of UPR via ceramide metabolism

As ceramide C20-C24 has been reported as a type of lipid to regulate UPR in cancer cells (*34, 35*), we employed a collection of assays to test the changes of UPR related factors in the adipose tissues or primary adipocytes. Initially, we observed the downregulation of genes related to the UPR by *Ntsr2* AKO in the WAT but not BAT (**Figure 4A-B and Supplementary Figure 6A**). *Ntsr2* AKO also downregulated the UPR associated proteins, including phosphorylated PERK (p-PERK) (*51, 52*) and phosphorylated eIF2α (p-eIF2α), in the WAT but not BAT (**Figure 4C-F**). RNA-seq analysis of WAT further revealed that the depletion of *Ntsr2* downregulated the expression of genes related to UPR in the WAT (**Supplementary Figure 6B**). In contrast, the phosphorylation of other integrated stress response factors, such as amino acid sensor GCN2 (*53–55*) and innate immunity response factor PKR (*56, 57*), were not changed by *Ntsr2* AKO (**Supplementary Figure 6C-D**). In addition, *Ntsr2* AKO did not change the expression of other upstream factors for UPR, including *Xbp1s* and *Atf6* (**Supplementary Figure 6E**). Interestingly, treatment of NTS peptide increased the level of UPR-related factors *in vivo* and *ex vivo* (**Figure 4G and Supplementary Figure 6F**).

**Figure 4.**
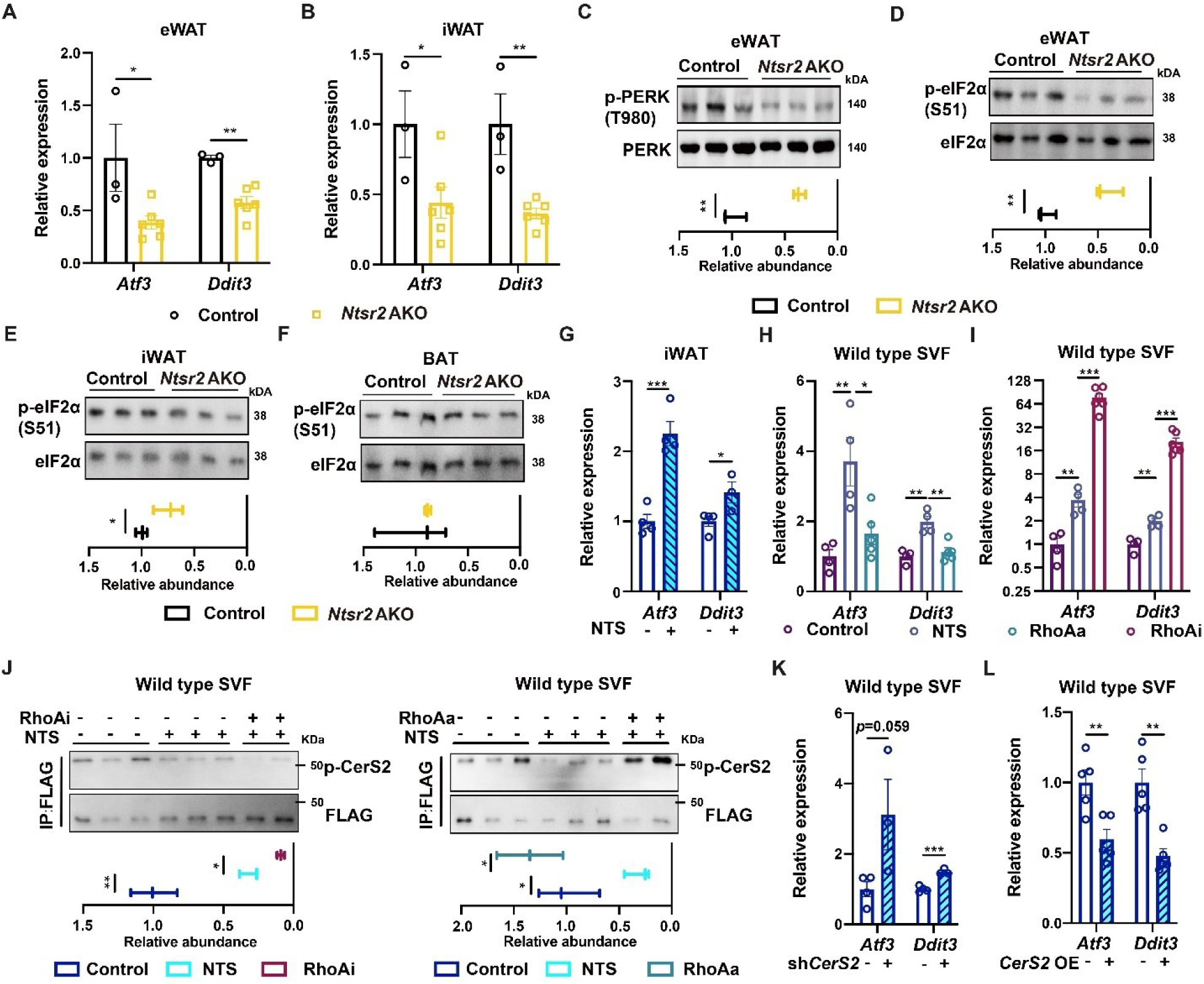
NTS-NTSR2 signaling regulated UPR via CerS2. **A-F.** The level of UPR related genes (**A-B**) and proteins (**C-F**, N=3) in the adipose tissues of control and *Ntsr2* AKO mice. N=3-6; **G.** The expression of UPR related genes upon NTS treatment *in vivo*. N=4; **H-I.** The expression of UPR related genes in the primary adipocytes upon the combinational treatment of NTS and RhoA agonist (RhoAa, **H**) or RhoA antagonist (RhoAi, **I**). N=4; **J.** The level of p-CerS2 in the primary adipocytes upon the combinational treatment of NTS and RhoAa (**left**) or RhoAi (**right**). N=2-3; **K-L.** The expression of UPR related genes in the primary adipocytes upon the knocking-down (**K**) or overexpression of *CerS2* (**L**). N=4-5. *, p<0.05; **, p<0.01; ***, p<0.001.

To further dissect the signaling between NTS-NTSR2 and UPR, we tested a collection of 13 chemicals which targeted various downstream factors of GPCR. Though most of the chemicals has no rational effects on the regulation of UPR-related genes by NTS, we did identify that RhoA was a potential factor linking the NTS-NTSR2 signliang and UPR in the primary adipcoytes. RhoA agonist (RhoAa) prevented the NTS treatment induced elevation of *Atf3* and *Ddit3* in the primary adipocytes (**Figure 4H**), but RhoA antagonist (RhoAi) enhanced the effects of NTS (**Figure 4I**). RhoAa treatment also prevented the decreases of phosphorylated CerS2 induced by NTS treatment, but RhoAi enhanced it (**Figure 4J**). These results together showed that NTS-NTSR2 signaling regulated the function of CerS2 via the RhoA.

Consistent to our hypothesis that CerS2 was a downstream factor of NTS-NTSR2 signaling, we discovered that the knockdown or overexpression of *CerS2* (**Supplementary Figure 7A-B**) controlled the expression of UPR-related factors (**Figure 4K-L and Supplementary Figure 7C**) in the primary adipocytes. Notably, the knocking-down of *CerS2* also increased the level of UPR-related factors in the primary adipocytes with *Ntsr2* depletion (**Supplementary Figure 7D**) and prevented the NTS-induced elevation of UPR-related factors (**Supplementary Figure 7E**). These data indicated that CerS2 is a downstream factor of NTS-NTSR2 signaling for regulating UPR. We also found the treatment of C22 ceramide, but not C16, downregulated the expression of UPR-related factors in the primary adipocytes (**Supplementary Figure 7F-H**). The *CerS2* KO mice presented the elevation of UPR-related factors in the WAT (**Supplementary Figure 7I-J**), but the local treatment of ceramide C22 *in vivo* downregulated the level of UPR-related factors in the WAT (**Supplementary Figure 7K**). These results suggested that NTS-NTSR2 signaling regulates the synthesis of ceramide C20-C24, which in turn affected the level of UPR in the WAT.

### NTS-NTSR2 signaling affected food intake via the downregulation of GDF15

The data derived from the *Ntsr2* AKO mice and the literature (*36*) inspired us to hypothesize that UPR, as a part of integrative stress response (ISR), may regulated the food intake via controlling the production of GDF15. Notably, we discovered that *Gdf15* gene, and its encoding product GDF15, were downregulated in the eWAT and iWAT (but not BAT) of *Ntsr2* AKO mice (**Figure 5A and Supplementary Figure 8A**). The GDF15 protein was also downregulated in the serum of *Ntsr2* AKO mice fed by either a chow diet (**Figure 5B**) or HFD (**Figure 5C**). In contrast, the local release of NTS peptide by hydrogel in the iWAT upregulated the serum GDF15 in both lean and obese mice (**Figure 5D and Supplementary Figure 8B**).

**Figure 5.**
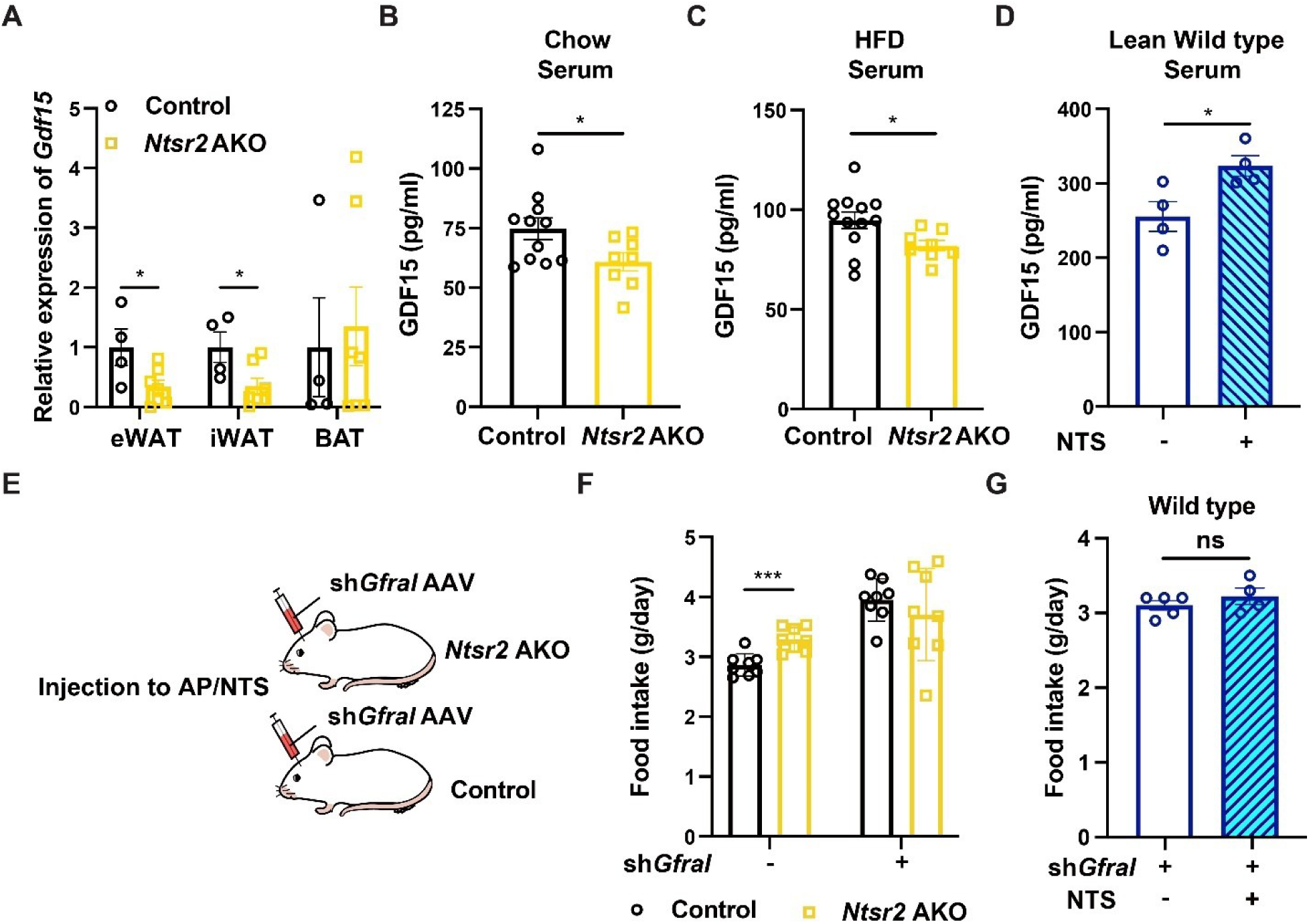
The decreases of GDF15 led to the increases of food intake in *Ntsr2* AKO mice. **A.** The expression of *Gdf15* genes in the adipose tissues; N=4-7; **B-C.** The abundance of GDF15 protein in the serum of control or *Ntsr2* AKO mice fed by a chow diet (**B**, N=8-11), HFD (**C**, N=8-12) or treated by NTS (**D**, N=5); **E.** The illustration of experimental design; **F.** The food intake of control or *Ntsr2* AKO mice with or without *Gfral* knocking-down. N=8; **G.** The food intake of mice treated by NTS in the iWAT locally with *Gfral* knocking-down. N=5. *, p<0.05; ***, p<0.001.

To validate whether the food intake differences between control and *Ntsr2* AKO mice were dependent on GDF15 signaling, we knocked down GDF15’s receptor *Gfral* (*58*) in the AP/NTS region of the brain via adeno-associated virus (AAV) injection (**Figure 5E**). As the interaction between GDF15 and GFRAL was well-established, we used it as a tool to validate our hypothesis. The effects of knocking-down on the expression of *Gfral* in the AP/NTS region were validated by fluorescent in situ hybridization (FISH) (**Supplementary Figure 8C**). The downregulation of *Gfral* also led to increases of food intake (**Supplementary Figure 8D**), which was consistent what has been reported. Notably, the changes of food intake induced by *Ntsr2* AKO or NTS local treatment in the iWAT was abolished by knocking down *Gfral* (**Figure 5F-G**). Simultaneously, *Ntsr2* AKO mice with *Gfral* knock-down were gaining body weight slower than the control mice when fed by a HFD (**Supplementary Figure 8E**). We also treated the mice with recombinant GDF15 via i.p. injection. We found that by normalizing the serum concentration of GDF15 (**Supplementary Figure 8F**), we managed to normalize the food intake of the control and *Ntsr2* AKO mice (**Supplementary Figure 8G**). In addition, we tried to normalize the serum concentration of GDF15 by enhancing its endogenous production via treating the mice with PERK agonist (PERKa) (*59*). Consistently, the PERKa treatment normalized the serum level of GDF15 as well as food intake in the *Ntsr2* AKO mice (**Supplementary Figure 8H-I**). These results indicated that NTS-NTSR2 signaling in the WAT regulates the food intake via GDF15 signaling. Notably, We recently have validated that the UPR related factor PERK controlled the food intake as well as metabolic homeostasis in the DIO mice via GDF15 (*60*).

We then tested whether the NTS-NTSR2 signaling regulated the production of GDF15 via controlling ceramide metabolism. We found that NTS treatment upregulated the expression of *Gdf15* in the SVF derived primary adipocytes comparing to the vehicle treatment, but this phenotype was abolished in the cells with the knocking-down of *CerS2* (**Figure 6A and Supplementary Figure 9A**) or with the treatment of RhoAa (**Supplementary Figure 9B**). The knocking-down of *CerS2* upregulated the production of GDF15 protein in both control and *Ntsr2* AKO primary adipocytes (**Figure 6B**). In contrast, the overexpression of *CerS2* downregulated production of GDF15 protein (**Figure 6C and Supplementary Figure 9C**) both control and *Ntsr2* AKO primary adipocytes. In addition, the treatment of C22 ceramide, but not C16, indeed downregulated the expression of *Gdf15* in the primary adipocytes (**Figure 6D and Supplementary Figure 9D**). The *CerS2* KO mice presented elevation of serum GDF15 concentration (**Figure 6E**), upregulation of *Gdf15* gene (**Figure 6F**) and GDF15 protein (**Figure 6G**) in the WAT comparing to the control mice. But the treatment of ceramide C22 *in vivo* downregulated the level of *Gdf15* gene or GDF15 protein in the WAT (**Supplementary Figure 9E**) and GDF15 protein in the serum (**Supplementary Figure 9F**). Together, these data suggested that NTS-NTSR2 signaling regulated the ceramide C20-C24 synthesis, which affected the production of GDF15 and food intake in mice.

**Figure 6.**
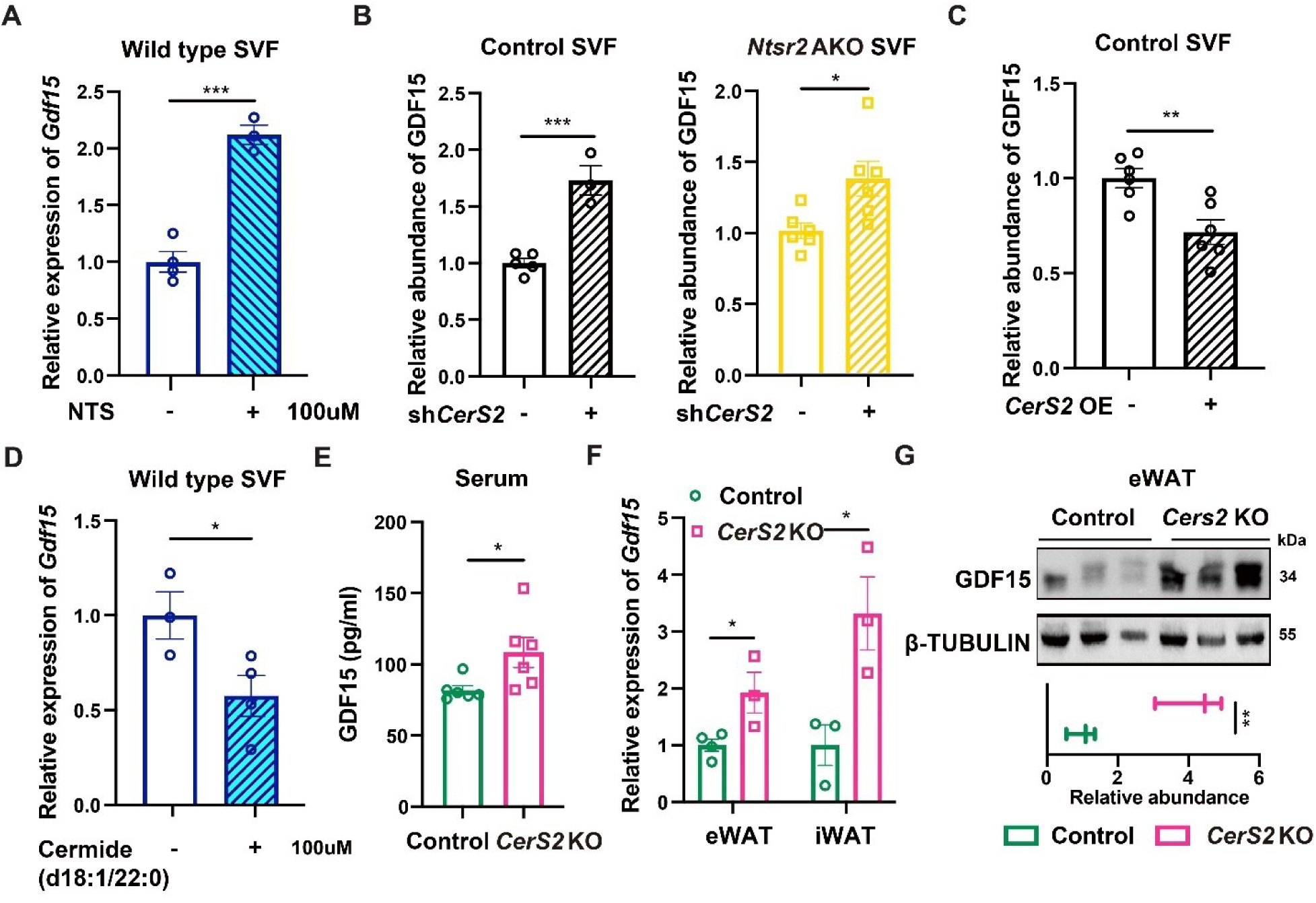
NTS-NTSR2 signaling regulated the production of GDF15 via CerS2. **A.** The relative expression of *Gdf15* upon NTS treatment in the primary adipocytes of wild-type mice; N=3-4; **B.** The relative production of GDF15 protein upon *CerS2* knock-down (**B**, N=3-5), overexpression (**C**, N=6) and ceramide C22 treatment (**D**, N=3-4) in primary adipocytes; N=3-6; **E-G.** The serum concentration of GDF15 (**E**, N=6), expression of *Gdf15* (**F**, N=3) and GDF15 protein abundance in adipose tissues (**G**, N=3) of control or *CerS2* KO mice. *, p<0.05; **, p<0.01; ***, p<0.001.

### The correlation between ceramides and food intake in human

To examine the generalization of our findings in human, we collected the relevant information and analyzed the correlation between plasma ceramide C20-C24 levels and daily intakes of food or energy in both children (282 participants) and adults (803 participants). The basic characteristics of these two populations were presented in **Figure 7A-B**. The positive associations between plasma levels of ceramide C20-C24 and energy intake were observed in both populations (**Figure 7C-F**). Based on the results from a recent systemic analysis for the correlation between serum concentration of metabolites and food intake, the correlation identified in the current study was among the top tier (*61*). Notably, a potential causal relation between plasma levels of ceramide C20-C24 and weight of daily food intake was observed using mendelian randomization analysis based on the data from UK Biobank and METSIM study (**Figure 7G; the technical details of mendelian randomization analysis were provided in the *“METHODS”* section**). These results suggested that the ceramide metabolism axis may also be of significance for the food intake in human.

**Figure 7.**
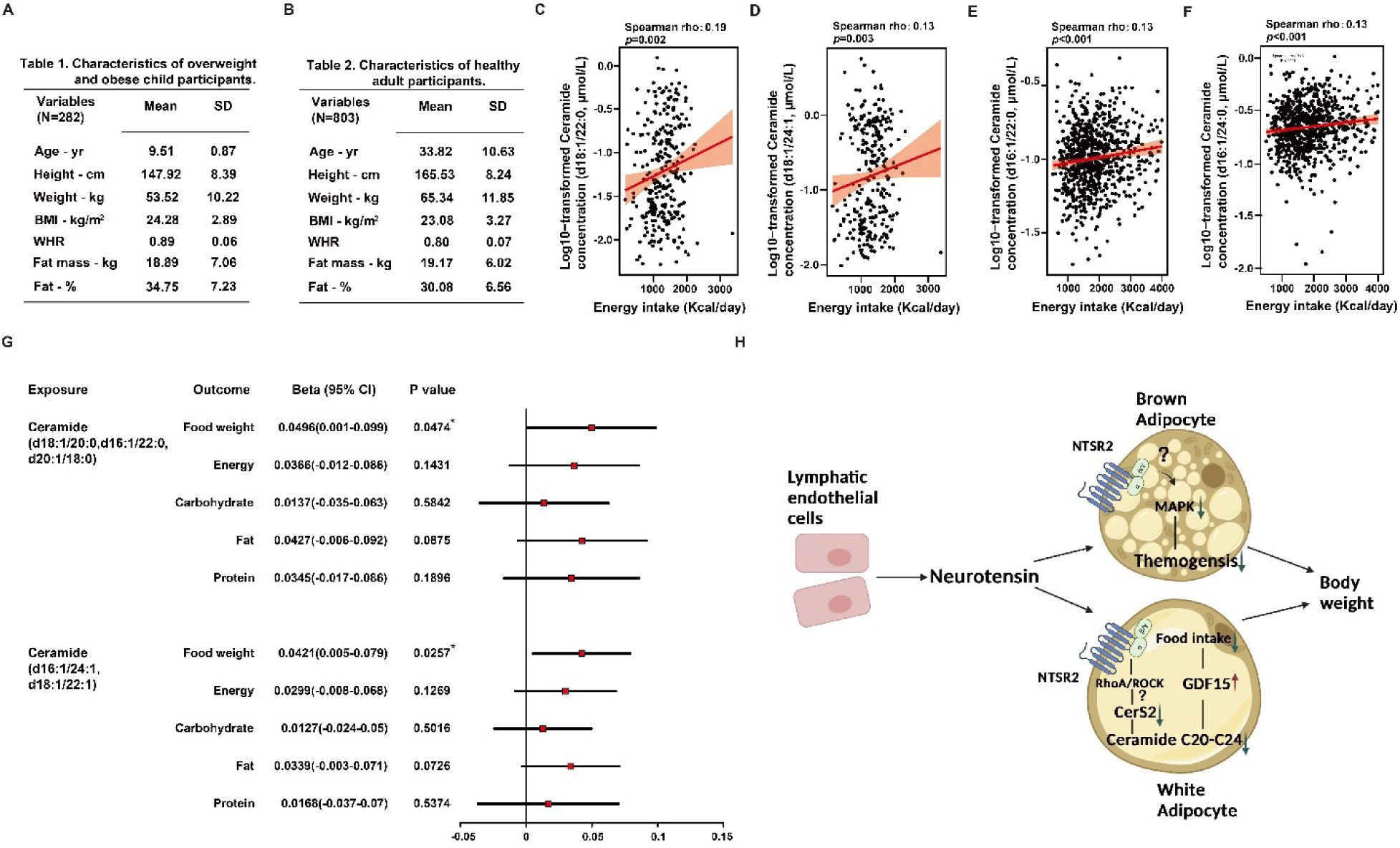
The relation between ceramide C20-C24 and food intake in human. **A-B.** The basic information of the children (**A**) and adult (**B**) populations; **C-F.** The correlation based on multivariable linear regression models between the serum concentration of ceramide C20-C24 and energy intake in the cohort of children (**C-D**) and adults (**E-F**); **G.** The summary of mendelian randomization analysis in adults. **H.** The illustration of the mechanism identified in this study.

## DISCUSSION

NTS has been recognized as an important peptide in the regulation of metabolic homeostasis for decades, though most studies on the functions and underlying mechanism of NTS were performed in the CNS or intestine. Our previous study has revealed the anti-thermogenic effects of LEC produced NTS via its interactions with NTSR2 in the BAT through the paracrine signaling. In the current study, we validated this thermogenic phenotype and further identified that NTS-NTSR2 signaling in the WAT regulated the food intake via its impacts on ceramide metabolism. NTS-NTSR2 signaling affected the synthesis of ceramide C20-C24, which induced the changes of GDF15 and food intake. These results indicated that the NTS-NTSR2 signaling in the peripheral tissues may play a key role in keeping the balance between BAT-dependent thermogenesis and WAT-dependent food intake. As the NTS-NTSR2 signaling can suppress thermogenesis in the BAT and suppress food intake in the WAT simultaneously, it is an important factor to maintain the metabolic homeostasis. The disruption of NTS-NTSR2 signaling in either of the tissues may lead to abnormal elevation of body weight as well as metabolic disorders (**Figure 7H**).

NTS can interact with both NTSR1 and NTSR2, but the downstream signaling pathways of these two GPCRs are strikingly different. Unlike NTSR1 (*62–66*), the effects of NTS stimulation on NTSR2 is controversial. For mouse NTSR2, NTS induced coupling of Gq to NTSR2 in a system of *Xenopus Laevis* oocytes but not HEK293 cells or COS cells (*67–70*). For human NTSR2, NTS behaved as a neutral antagonist in COS cells and HEK293 cells but was an activator of MEK-ERK signaling in CHO cells (*71–74*). Previously, we have reported that NTS controlled the MEK-ERK signaling in the BAT. But the results of current study indicated that the MEK-ERK signaling was not affected by the depletion of *Ntsr2* in the WAT. The tissue and context-specificity are the outstanding features for the function of GPCR (*75*). These studies with contradictory conclusions highlighted the importance of studying NTS-NTSR2 signaling with cell type specific models in physiological conditions.

In an *ex vivo* model, we have previously identified that NTS activates MEK-ERK signaling which is responsible for its anti-thermogenic effects (*23*). In the current study, we discovered that NTS may exert inhibitory effects on food intake via NTSR2 in WAT, but not BAT, via regulating the RhoA-CERS2-GDF15 axis *in vivo*. As far as we know, this is the first evidence to support the hypothesis that NTS may exert neurological regulation via direct interactions with peripheral tissues. Further studies are required to elucidate whether it a common mechanism or adipocyte specific phenomena.

Previously, we have reported that treatment with the NTSR2 antagonist NTRC-824 enhanced thermogenesis without obvious effects on food intake in mice (*23*). Encouragingly, the *Ntsr2* BKO and AKO mice in the current study also presented enhance thermogenesis. The inconsistencies in the food intake phenotypes between *Ntsr2* AKO mice and mice treated with NTRC-824 may be due to several differences between the two studies: **a)** NTRC-824 treatment exerted its effects on the whole body in comparison to adipocyte specific depletion of *Ntsr2* shown in the current study. The complicated effects of NTSR2 inhibition on various cell types may disturb the feeding behavior of mice; **b)** NTRC-824 treatment was performed on obese mice while *Ntsr2* depletion happened when mice were still lean. The dramatic differences between lean and obese mice may affect the phenotypes associated with the feeding behavior; **c)** Many chemicals have more than one target *in vivo*. Similarly, NTRC-824 may have unknown off-target effects other than inhibiting the activity of NTSR2.

Ceramide has been broadly recognized as a type of lipid with important bioactivities. Ceramide may exert diverse effects on metabolism related phenotypes via interactions with brown/beige adipocytes or the CNS (*76–78*). Many groups, including ourselves, have demonstrated that the total ceramide is an important factor for senescence or aging (*79, 80*). Notably, it has been reported that the intracerebroventricular (ICV) administration of mixed species of ceramide to rainbow trout induced decreases of food intake in fish (*81*). In contrast, the effects of ceramide on the food intake of mammals were complicated. ICV injection or local ablation of ceramide C6 in hypothalamus of mice may either increase food intake (*82*) or have no effects on food intake (*77*). The current studies identified that the ceramide C20-C24, as the specific products of CerS2 from peripheral tissues, may regulate the food intake in both mice and human. The metabolism of ceramide has the potential to become a therapeutic target for obesity.

The unfolded protein response is an important pathway for the regulation of endoplasmic reticulum (ER) stress. It has been reported in several studies that the intracellular abundance of ceramide was associated with ER stress (*83*). In the macrophage, the ASMase can induce ER stress via elevation of ceramide and oxidized-LDL (*84*). In the CNS, the treatment of ceramide synthesis inhibitor myriocin or depletion of *CerS6* suppressed the ER stress in the hypothalamus (*78, 85*). Especially, it has been discovered that the dihydroceramide can bind directly to ATF6, a key regulator of UPR, which activated the downstream signaling (*86*). We therefore hypothesized that the C20-24 ceramide may specifically interact with a protein involved in the regulation of PERK. This interaction controlled the phosphorylation of PERK, the production of GDF15 and the food intake. As the current study is a physiological study in general, this specific protein interacting with C20-24 ceramide was waiting to be discovered by biophysical/chemical biology studies with a separated project.

In summary, our study provided an inspiring insight into the function and relevant mechanism of NTS-NTSR2 signaling in the context of metabolic regulation. In addition to NTS, other secretory proteins produced by the LEC have been discovered (*87–91*). Together, these studies will inspire the future work on how the LEC interacts with peripheral organs/tissues via secretory proteins, as well as how it regulates metabolic homeostasis.

## METHODS

### Reagents

The information of key reagents involved in this study can be provided upon request.

### Mouse models

All animal experiments were performed according to procedures approved by the Animal Ethics Committee of Fudan University. Mice were maintained under a 12 hr light/12hr dark cycle at constant temperature (23°C) with free access to food and water. Wild-type C57B6/J mice, *Ntsr2^flox/flox^* mice and *Cers2*^+/-^ mice were obtained from GenPharmatech. The Ucp1-cre and Adipoq-cre mice were obtained from Jax Lab.

### Local release of NTS

The hydrogel was synthesized and the NTS peptide was loaded to the hydrogel as described before (*92, 93*). In brief, the sodium hyaluronate, poloxamer 407 and ultrapure water were mixed, centrifuged and store in the fridge. The NTS peptide were added to the hydrogel as 1.5mg/ml. For the treatment, 100 µl of hydrogel with or without NTS was injected to the iWAT once every two days. The tissues and serum were collected from mice with treatment for 6 days.

### Culture and differentiation of mouse pre-adipocytes

Inguinal adipose tissue was dissected from 7–10-week-old C57/Bl6J male mice, minced, and digested in PBS with 10 mg/ml BSA (Cat#A600332-0100, BBI Life Sciences) and 10 mg/mL collagenase (Cat#11088858001, Roche) for 60 min at 37°C. The digested solution was filtered through a 100 µm cell strainer, centrifuged at 800 rpm for 5 min and the red blood cells were lysed by ACK buffer (Cat# A10492-01, Gibco). The stromal-vascular fraction (SVF) pellet was then suspended in the DMEM medium with 10% FBS and 1% P/S for cell culture experiments. The SVF cells were differentiated into adipocyte with growth medium containing 5 ug/mL insulin, 0.5 mM isobutylmethylxanthine, 1 uM rosiglitazone, 1 uM dexamethasone for two days initially. Then we refed the cells with new growth medium containing 5 ug/mL insulin for another two days. Afterwards, the cells were maintained in growth medium without insulin. Cells were fully differentiated after 6-8 days (*94*). As the expression of *Ntsr2* in the differentiated adipocytes was not detectable, we overexpressed *Ntsr2* in the adipocytes via the infection of adeno virus. The sequences for shRNA can be provided upon request.

### Western blotting

The tissues were homogenized in 1% NP40 buffer (*95*) containing phosphatase inhibitor cocktail (Yeasen Biotech) and protease inhibitor cocktail (APE×BIO) with metal beads. The lysates were then separated by SDS-PAGE and transferred to Immobilon 0.45mm membranes (Cat# IPVH00010, Millipore). Western blotting was conducted using specific various primary and secondary antibodies. The membranes were incubated diluted primary antibodies overnight at 4°C and secondary antibodies for 1 hour at room temperature. ECL Reagent (Cat# G2014-50ML Servicebio) and ChemiScope machine (NO. 6000, Clinx Science Instruments Co., Ltd) was used for the visualization of protein bands.

### Serum ELISA

The serum samples were collected by centrifuging the whole blood of mice for 10 minutes at 4000 rpm and stored in a deep freezer. The levels of GDF15 protein in the serum were measured by an ELISA kit (Cat# MGD150, R&D Systems) following the manufacturer’s instructions.

### Human studies

All participants in both populations provided written informed consent. The cohort of children was recruited from two elementary schools in Beijing, China. This study has been approved by the ethics committee of Peking University Third Hospital (ethnic approval number 2021-283-06). The cohort was established base on the related protocol (*96*) . The information of 282 donors were used in this study. The cohort of adults recruited 803 healthy individuals free of diabetes, gastrointestinal disease, cardiovascular diseases, cancer, or any use of medication in Shanghai, China. This study has been approved by the Ethics Committees of Fudan University (FE20064) and Zhongshan Hospital (B2019-089R). The details of the two populations were described in the supplementary information. The samples were analyzed by untargeted lipidomics, which was described below.

### Tissue lipid extraction

Lipid extraction was performed following a published protocol (*97*). For serum samples, they were centrifuged to remove cells at 4°C, 14000× g for 10 min. 30 μL of serum, 120 μL of water, 1.5 mL of methanol and 5 mL of MTBE were added to a clean glass centrifuge tube, and 12 μL of lipid internal standards were added in each sample at the same time, including 2 μL of 0.82 mM PC(17:0/17:0), 2 μL of 0.77 mM PC(19:0/19:0), 2 μL of 1.40 mM PE (14:0/14:0), 2 μL of 4.91 mM LPC (17:0), 2 μL of 3.49 mM SM (d18:1/17:0) and 2 μL 4.53 mM of Cer (d18:1/17:0). The homogenate was vortexed for 1 min. The glass centrifuge tube containing the homogenate was rocked on a shaker for 1 h at room temperature. A total of 1.25 mL of water was then added to the glass centrifuge tube followed by another minute of vortexing. The homogenate was centrifuged at 4 °C, 1000× g for 10 min and two-phase layers could be observed in the glass centrifuge tube. A total of 4 mL of the top phase supernatant were collected and dried under a stream of nitrogen. The extracted lipid samples were stored at −80 °C before LC-MS/MS analysis. For the adipose tissues, 2 mg eWAT tissue sample was added to 200 µL of water and 500 µL of methanol and homogenized using the same approach as in the hydrophilic metabolite extraction above. The homogenate was supplemented with 500 µL more methanol and decanted into a clean glass centrifuge tube. Five milliliters of MTBE were then added to the glass centrifuge tube and vortexed for one min. The glass centrifuge tube containing the homogenate was rocked on a shaker for one hour at room temperature. 1.25 mL of water was then added to the glass centrifuge tube followed by another minute of vortexing. The homogenate was centrifuged at 4°C at 1,000xg for 10 min and two-phase layers could be observed in the glass centrifuge tube. Four milliliters of the top phase supernatant were collected and dried under a stream of nitrogen. The extracted lipid sample was stored at -80°C before MS analysis.

### Untargeted lipidomics

The untargeted lipidomics method was modified from a published method (*98*). Lipid samples were resuspended in 50 µL of isopropanol: acetonitrile: water (vol/vol/vol,30:65:5), and 10 µL was injected into Orbitrap Exploris 480 LC-MS/MS (Thermo, USA) coupled to HPLC system (Shimadzu, Kyoto, Japan). Lipids were eluted via C30 by using a 3 μm, 2.1 mm × 150 mm column (Waters) with a flow rate of 0.26 mL/min using buffer A (10 mM ammonium formate at a 60:40 ratio with acetonitrile: water) and buffer B (10 mM ammonium formate at a 90:10 ratio with isopropanol: acetonitrile). Gradients were held in 32% buffer B for 0.5 min and run from 32% buffer B to 45% buffer B at 0.5–4 min; from 45% buffer B to 52% buffer B at 4–5 min; from 52% buffer B to 58% buffer B at 5–8 min; from 58% buffer B to 66% buffer B at 8–11 min; from 66% buffer B to 70% buffer B at11–14 min; from 70% buffer B to 75% buffer B at 14–18 min; from 75% buffer B to 97% buffer B at 18–21 min; 97% buffer B was held from 21–25 min; from 97% buffer B to 32% buffer B at 25–25.01 min; and 32% buffer B was held for 8 min. All the ions were acquired by non-targeted MRM transitions associated with their predicted retention time in a positive and negative mode switching fashion. ESI voltage was +5500 and −4500V in positive or negative mode, respectively. All the lipidomics .RAW files were processed on *LipidSearch* 4.0 (Thermo Fisher Scientific) for the lipid identification. The lipidomic results were statistically analyzed with MetaboAnalyst 5.0 (*99*) and LINT-web (*100*).

### Targeted Lipidomic Analysis of Ceramide

Samples were resuspended in 100 μL of isopropanol: acetonitrile: water (v:v:v, 65:30:5), and 10 μL was injected into a 6500 QTRAP triple-quadrupole MS (SCIEX) coupled to an HPLC system (Shimadzu). Ceramides were eluted via C30 by using a 3 μm, 2.1 mm × 150 mm column (Waters) with a flow rate of 0.26 mL /min using buffer A (10 mM ammonium formate at a 60:40 ratio with acetonitrile: water) and buffer B (10 mM ammonium formate at a 90:10 ratio with isopropanol: acetonitrile). Gradients were held in 32% buffer B for 0.5 min and run from 32% buffer B to 45% buffer B at 0.5–4 min; from45% buffer B to 52% buffer B at 4–5 min; from 52% buffer B to 58% buffer B at 5–8 min; from 58% buffer B to 66% buffer B at 8–11 min; from 66% buffer B to 70% buffer B at 11–14 min; from 70% buffer B to 75% buffer B at 14–18 min; from 75% buffer B to 97% buffer B at 18–21 min; 97% buffer B was held from 21–25 min; from 97% buffer B to 32% buffer B at 25–25.01 min; and 32% buffer B was held for 8 min.

Ceramides have similar precursor ions, such as d18:1 sphingosine (m/z 264 and 282), d16:1 sphingosine (m/z 236 and 254) and t18:0 sphingosine (m/z 282 and 300) (*33*). All ions were acquired by 156 selected reaction monitoring transitions just in a positive mode. Electrospray ionization (ESI) voltage was +4900 V.

### Data representation and statistical analysis

The number of replicates were described in the figure legends. All the values in the figure are expressed as mean ± SEM. All the *p*-values were calculated via t-test with post-hoc correction unless specifically indicated.

## ACKNOWLEDGMENTS

We thank for the discussion with Dr. Evan Rosen (Harvard Medical School). We thank Single Cell Quantitative Metabolomics and Lipidomics Core Facility of IMIB at Fudan University for LC-MS/MS analysis. This work was supported by MOST 2020YFA0803800, 2019YFA0801900, NSFC92057115, and Shanghai Sailing Program 20YF1402600 to H.H.; MOST 2020YFA0803600, 2018YFA0801300, 2023YFA1801000, NSFC 32071138, 32371193 and SKLGE-2118 to J.L.

## AUTHOR CONTRIBUTIONS

Conceptualization, W.F., J.L.; Investigation, W.F., Y.Y., X.G., Q.G., X.Z., L.Z., C.L., Z.Z., J.S., Y.Z., L.H., G.L., C.Y., Q.W., C.F., C.L., J.W., H.Y., Y.Z., T.H., H.R., T.L., H.T., J.Q., Y.Z., M.X., H.H., J.L.; Analysis, W.F., Y.Y., X.G., Q.G., X.Z., L.Z., C.L., Z.Z., J.S., Y.Z., L.H., G.L., C.Y., Q.W., C.F., C.L., J.W., H.Y., Y.Z., T.H., H.R., T.L., H.T., J.Q., Y.Z., M.X., H.H., J.L.; Writing, W.F., Y.Y., X.G., Q.G., X.Z., L.Z., C.L., Z.Z., J.S., Y.Z., L.H., G.L., C.Y., Q.W., C.F., C.L., J.W., H.Y., Y.Z., T.H., H.R., T.L., H.T., J.Q., Y.Z., M.X., H.H., J.L.; Data Visualization, W.F., Y.Y., X.G., Q.G., X.Z., L.Z., C.L., Z.Z., J.S., Y.Z., L.H., G.L., C.Y., Q.W., C.F., C.L., J.W., H.Y., Y.Z., T.H., H.R., T.L., H.T., J.Q., Y.Z., M.X., H.H., J.L.; Funding Acquisition, J.L.; Supervision, Y.Z., H.H., J.L..

---

**Supplementary Figure 1.**
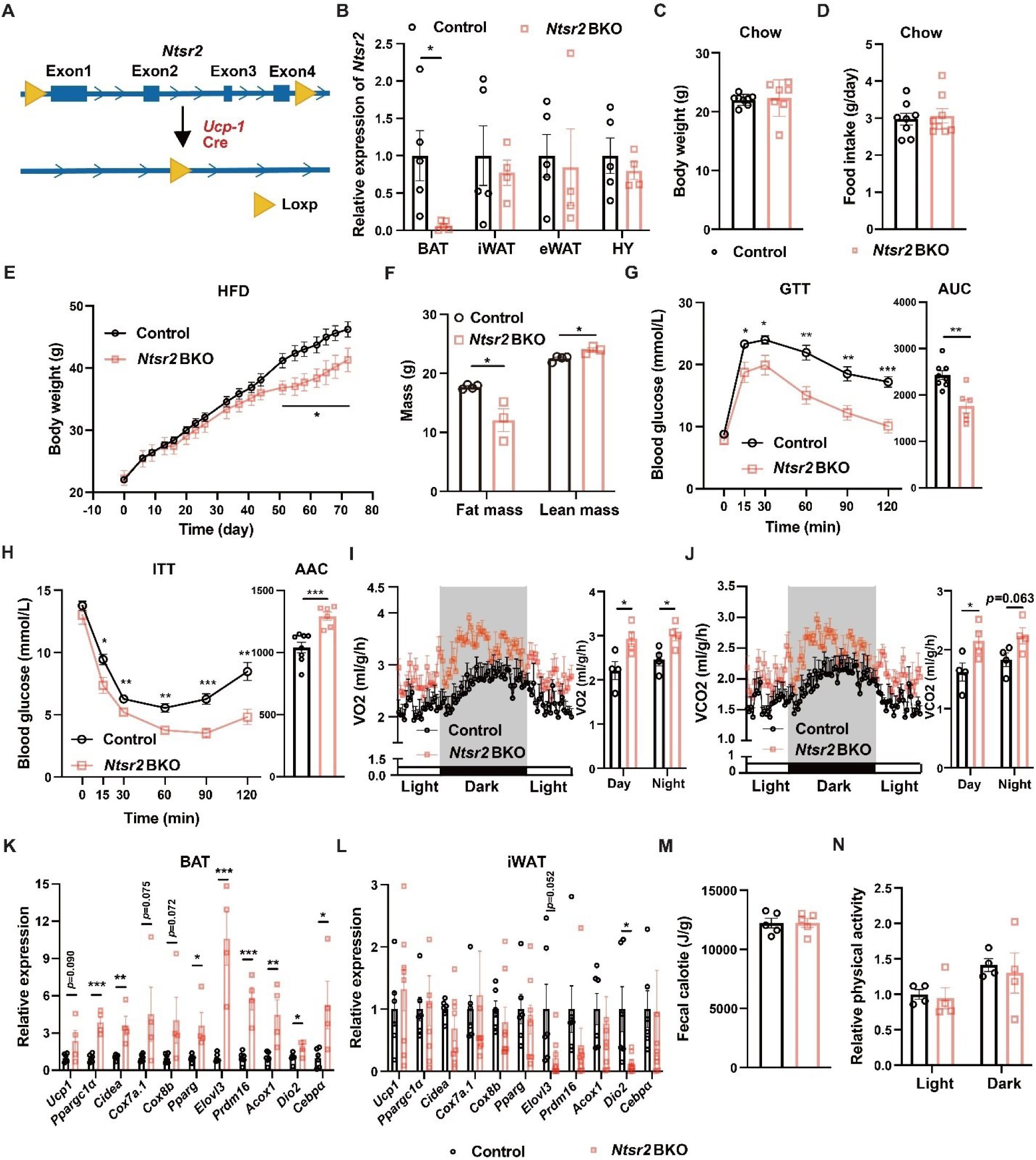
The elevation of thermogenesis in *Ntsr2* BKO mice. **A.** Establishment of brown/beige adipocyte specific *Ntsr2* KO mouse model; **B.** The KO efficiency and specificity of the *Ntsr2* gene; N=4-5; **C-D.** The body weight (**C**) and food intake (**D**) of mice fed by a chow diet. N=7; **E-N.** The body weight (**E**), body composition (**F**), GTT (**G**), ITT (**H**), O2 (**I**) or CO2 (**J**) production rate, the expression of thermogenic genes in BAT (**K**) or iWAT (**L**), fecal energy (**M**) and physical activity (**N**) of mice fed by HFD. N=7-8. *, p<0.05; **, p<0.01; ***, p<0.001. HY, hypothalamus; iWAT, inguinal white adipose tissue; eWAT, epidydimal white adipose tissue; BAT, brown adipose tissue.

**Supplementary Figure 2.**
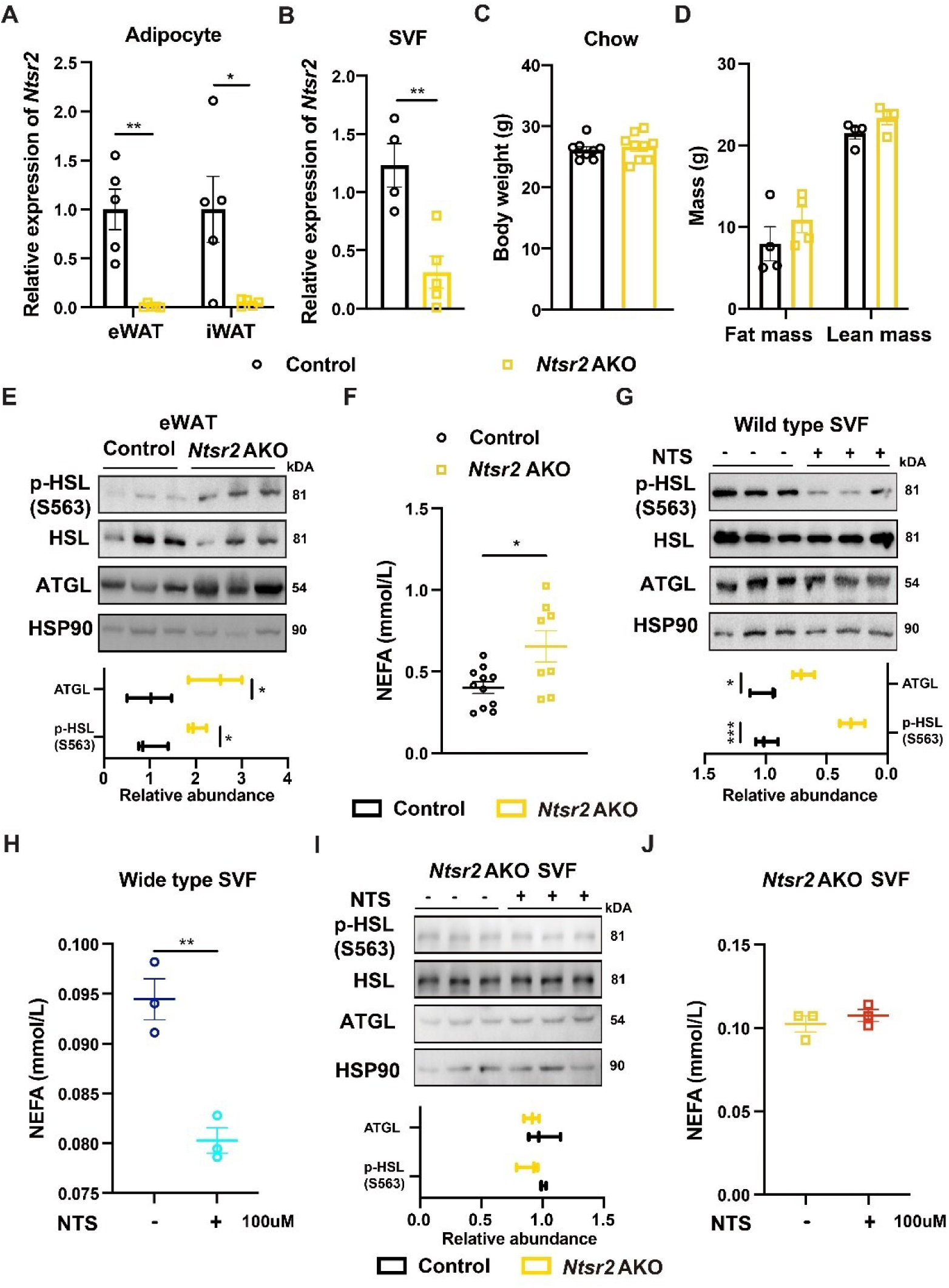
NTS-NTSR2 signaling inhibits the lipolysis of WAT. **A.** The KO efficiency in the floated adipocytes. N=5; **B.** The KO efficiency in the primary adipocytes. N=4-5; **C.** The body weight (**C**) and body composition (**D**) of mice fed by chow diet; **E.** The expression of p-HSL and ATGL protein in the eWAT; **F.** The concentration of NEFA in the serum of control or *Ntsr2* AKO mice. N=8-10; **G-H.** The expression of p-HSL and ATGL protein (**G**) as well as NEFA secretion (**H**) from the primary adipocytes of the wild type mice. N=3; **I-J.** The expression of p-HSL and ATGL protein (**I**) as well as NEFA secretion (**J**) from the primary adipocytes of the *Ntsr2* AKO mice. N=3. *, p<0.05; **, p<0.01; ***, p<0.001.

**Supplementary Figure 3.**
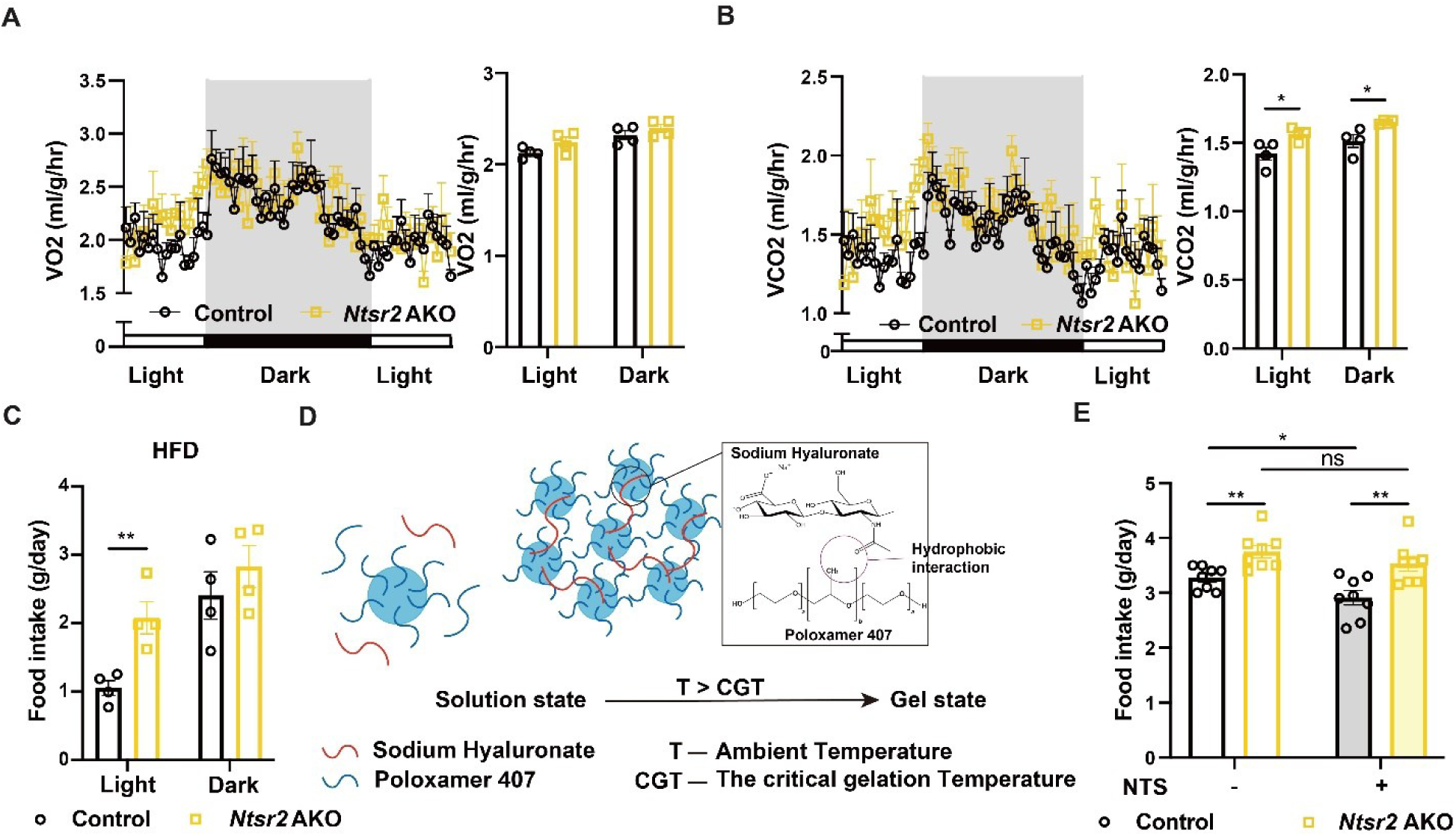
The impacts of NTS-NTSR2 signaling on metabolic homeostasis. **A-B.** The O2 consumption rate (**A**) and CO2 production rate (**B**) of the mice fed by HFD; N=4. **C.** The food intake of control and *Ntsr2* AKO mice fed by an HFD. N=8; **D.** The illustration of hydrogel containing NTS peptide; **E.** The effects of NTS local release on the food intake of control and *Ntsr2* AKO mice. N=8. *, p<0.05; **, p<0.01.

**Supplementary Figure 4.**
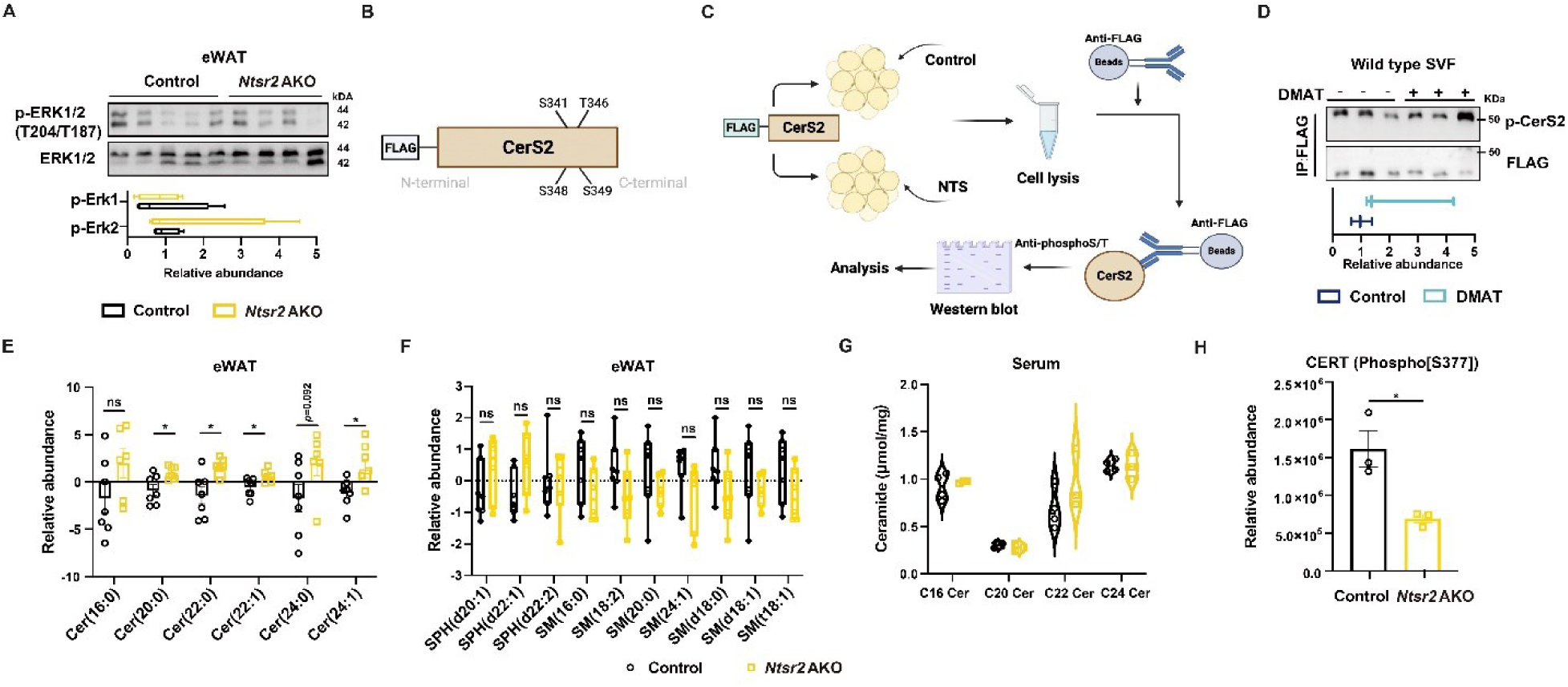
The metabolism of ceramide controlled by the NTS-NTSR2 signaling in the WAT. **A.** The level of phosphorylated ERK (p-ERK) in the WAT of control and *Ntsr2* AKO mice; **B.** The illustration of phosphorylation site for CerS2; **C.** The illustration of how to detect the phosphorylated CerS2 (p-CerS2) with the pull-down and western blot assay; **D.** The level of p-CerS2 with the treatment of CK2 inhibitor DMAT; **E-F.** The relative abundance of ceramide (**E**) or sphingosine (**F**) in the WAT, detected by the untargeted lipidomics. N=5-6; **G.** The concentration of ceramide in the serum of control and *Ntsr2* AKO mice. N=5-6; **H.** The relative abundance of phosphorylated CERT (p-CERT) in the WAT. N=3. *, p<0.05.

**Supplementary Figure 5.**
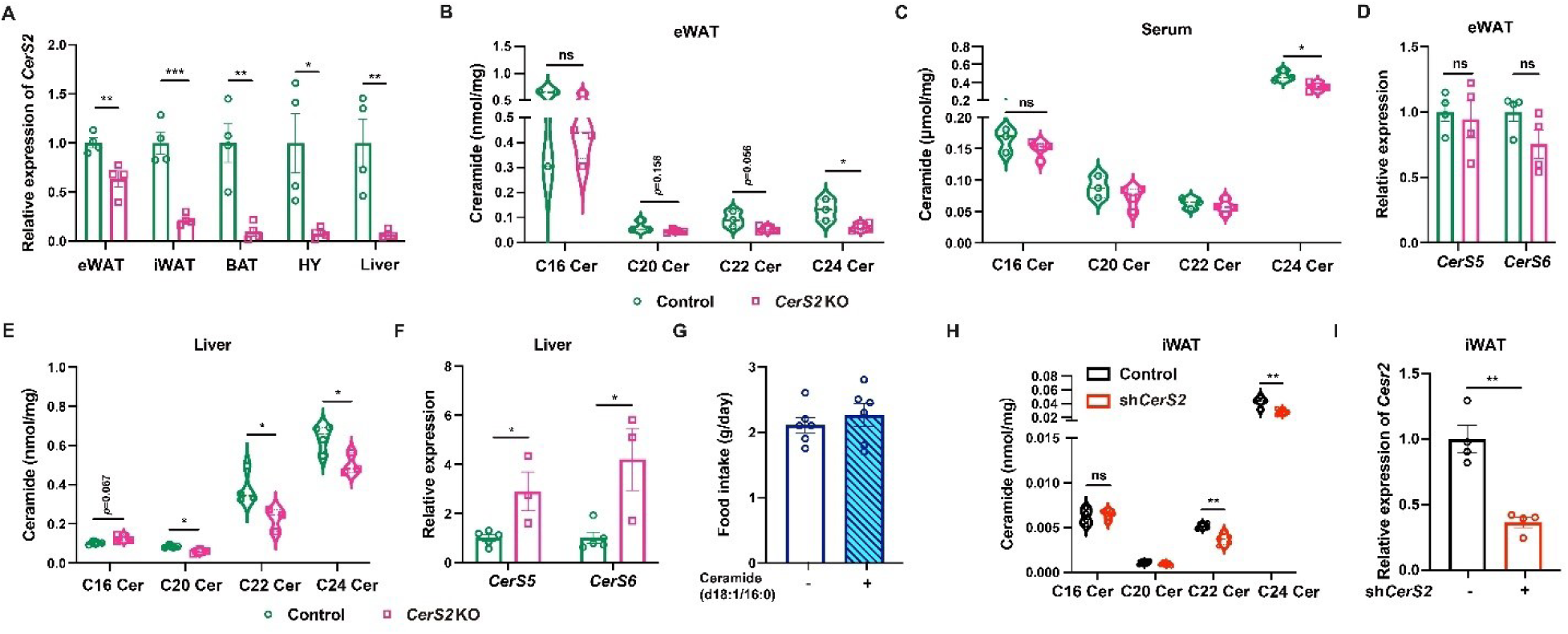
CerS2 in the WAT regulated the metabolic homeostasis. **A.** The KO efficiency of *CerS2* detected by RT-qPCR. N=4; **B-C.** The concentration of ceramide C16-24 in the WAT (**B**) or serum (**C**) of control or *CerS2* KO mice. N=4; **D.** The expression of *CerS5* and *CerS6* in the WAT. N=4; **E-F.** The concentration of ceramide C16-24 (**E**) as well as the expression of *CerS5* and *CerS6* (**F**) in the liver of control or *CerS2* KO mice. N=4; **G.** The food intake of mice treated with ceramide C16. N=6; **H-I.** The concentration of ceramide C16-24 (**H**) and the expression of *CerS2* (**I**) with the knocking-down of *CerS2* in the iWAT locally. N=4. *, p<0.05; **, p<0.01; ***, p<0.001.

**Supplementary Figure 6.**
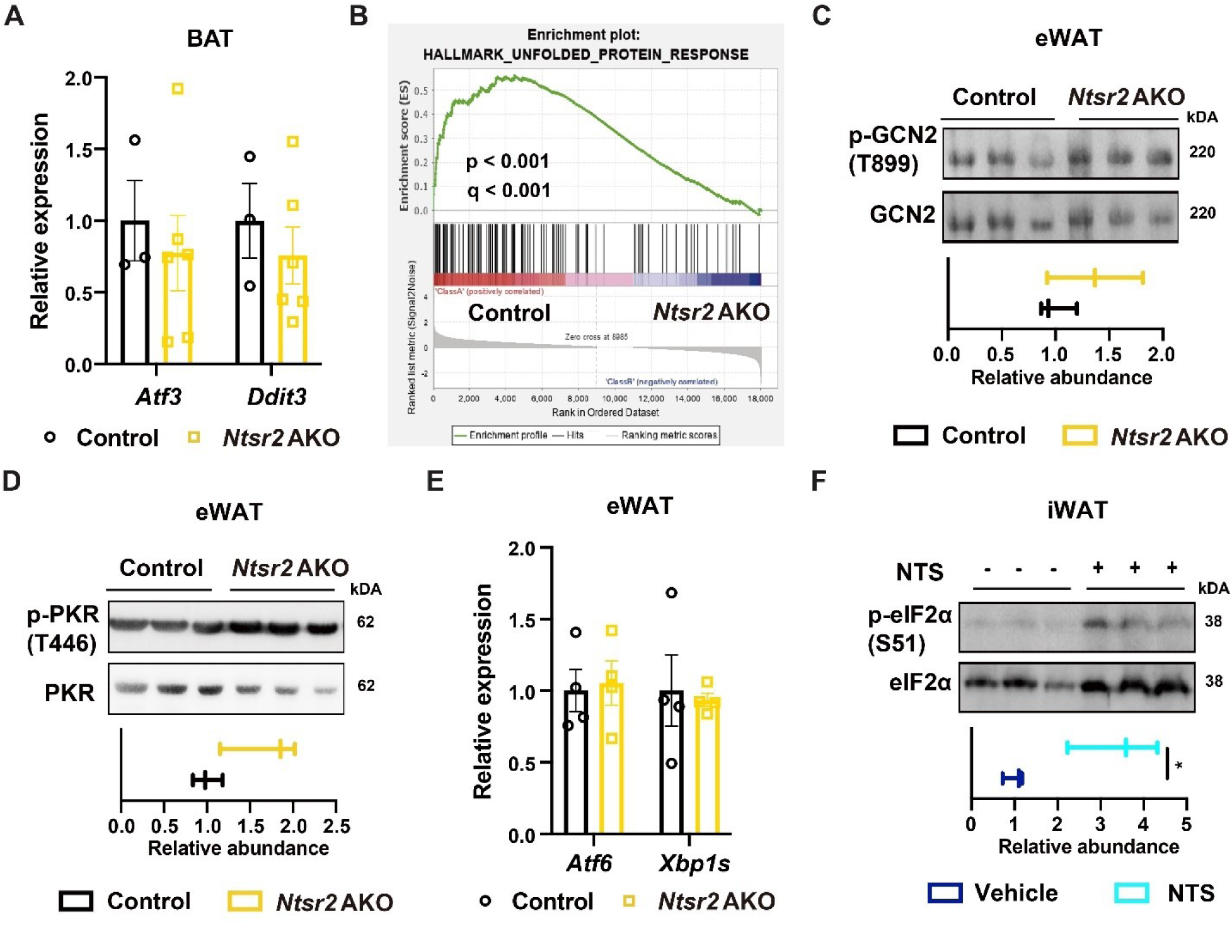
NTS-NTSR2 signaling regulated UPR. **A.** The expression of UPR related genes in BAT. N=3-6; **B.** GSEA revealed the downregulation of UPR related genes in the WAT of control and *Ntsr2* AKO mice. N=3; **C-D.** The level of p-GCN2 (**C**) and p-PKR (**D**) in the WAT of control and *Ntsr2* AKO mice. N=3; **E.** The expression of two UPR related genes in the WAT of control and *Ntsr2* AKO mice. N=4; **F.** The level of p-eIF2α in the primary adipocytes upon NTS treatment. N=3. *, p<0.05.

**Supplementary Figure 7.**
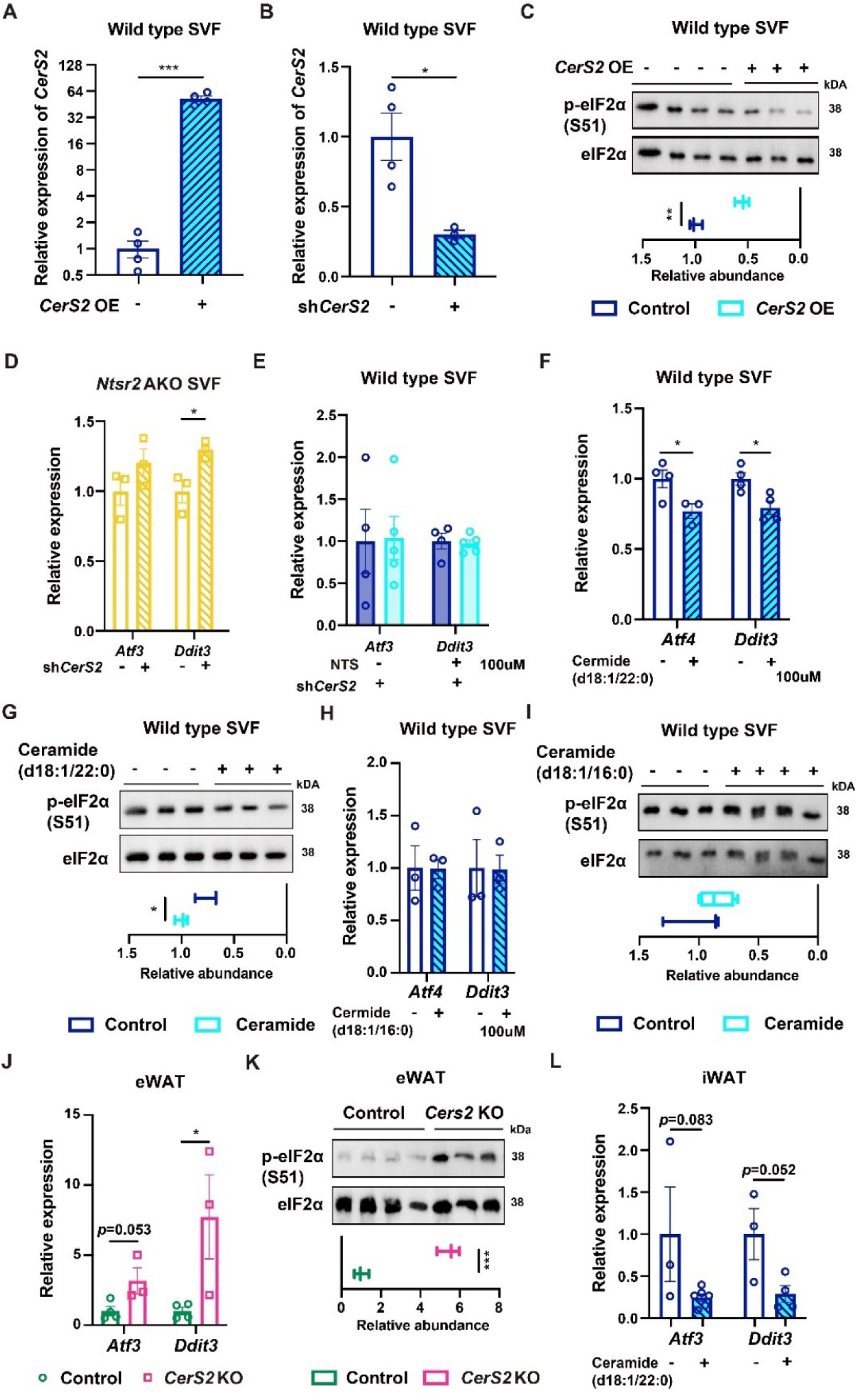
Ceramide metabolism regulated UPR. **A-B.** The validation of overexpression (OE, **A**) and knocking-down (**B**) of *CerS2* in the primary adipocytes. N=4; **C.** The level of p-eIF2α in the primary adipocytes upon the overexpression of *CerS2*. N=3-4; **D.** The expression of UPR related genes in the primary adipocytes with *Ntsr2* AKO and knocking-down of *CerS2*. N=3; **E.** The expression of UPR related genes in the primary adipocytes with knock-down of *CerS2* upon NTS treatment. N=4-5; **F-G.** The expression of UPR related genes (**F**) and p-eIF2α (**G**) in the primary adipocytes upon ceramide C22 treatment. N=3-4; **H-I.** The expression of UPR related genes (**H**) and p-eIF2α (**I**) in the primary adipocytes upon ceramide C16 treatment. N=3-4; **J-K.** The expression of UPR related genes (**J**) and p-eIF2α (**K**) in the WAT of control or *CerS2* KO mice. N=3; **L.** The expression of UPR related genes in the WAT upon ceramide C22 *in vivo*. N=3-4. *, p<0.05; ***, p<0.001.

**Supplementary Figure 8.**
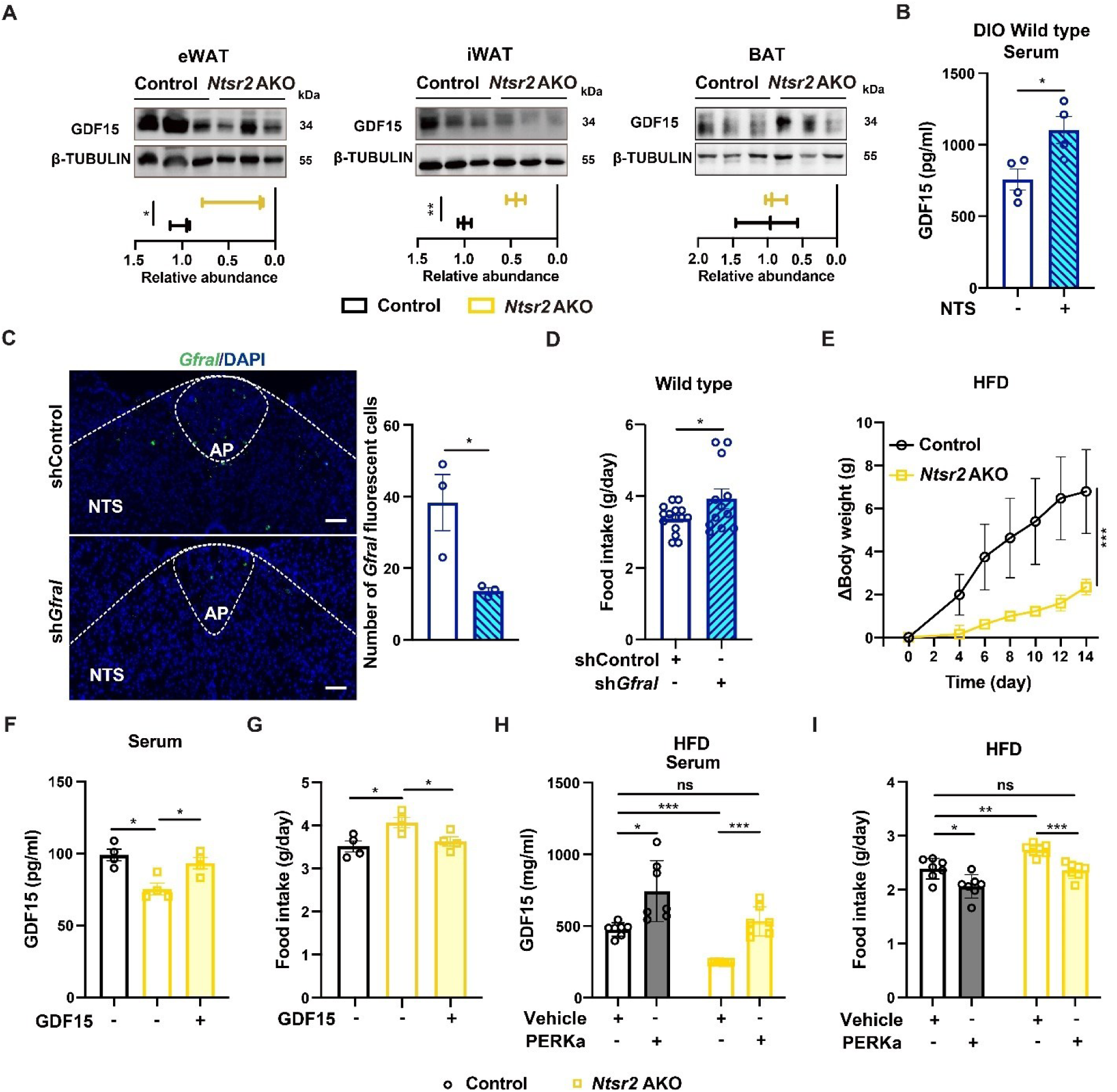
NTSR2 regulated food intake via GDF15. **A.** The abundance of GDF15 protein in the adipose tissue. N=3; **B.** The serum concentration of GDF15 from the obese mice treated with NTS in the iWAT locally. N=4; **C.** The expression of *Gfral* in the AP/NTS region with or without *Gfral* knock-down. N=3; **D.** The food intake of mice with or without *Gfral* knock-down; N=12-14; **E.** The body weight changes of control or *Ntsr2* AKO mice with or without *Gfral* knock-down; N=3; **F-G.** The recombinant GDF15 treatment normalized the serum concentration of GDF15 (**F**) and food intake (**G**) in the control or *Ntsr2* AKO mice. N=4; **H-I.** The treatment of PERK agonist (PERKa) normalized the serum concentration of GDF15 (**H**) and food intake (**I**) in the control and *Ntsr2* AKO mice. N=7. *, p<0.05; **, p<0.01; ***, p<0.001.

**Supplementary Figure 9.**
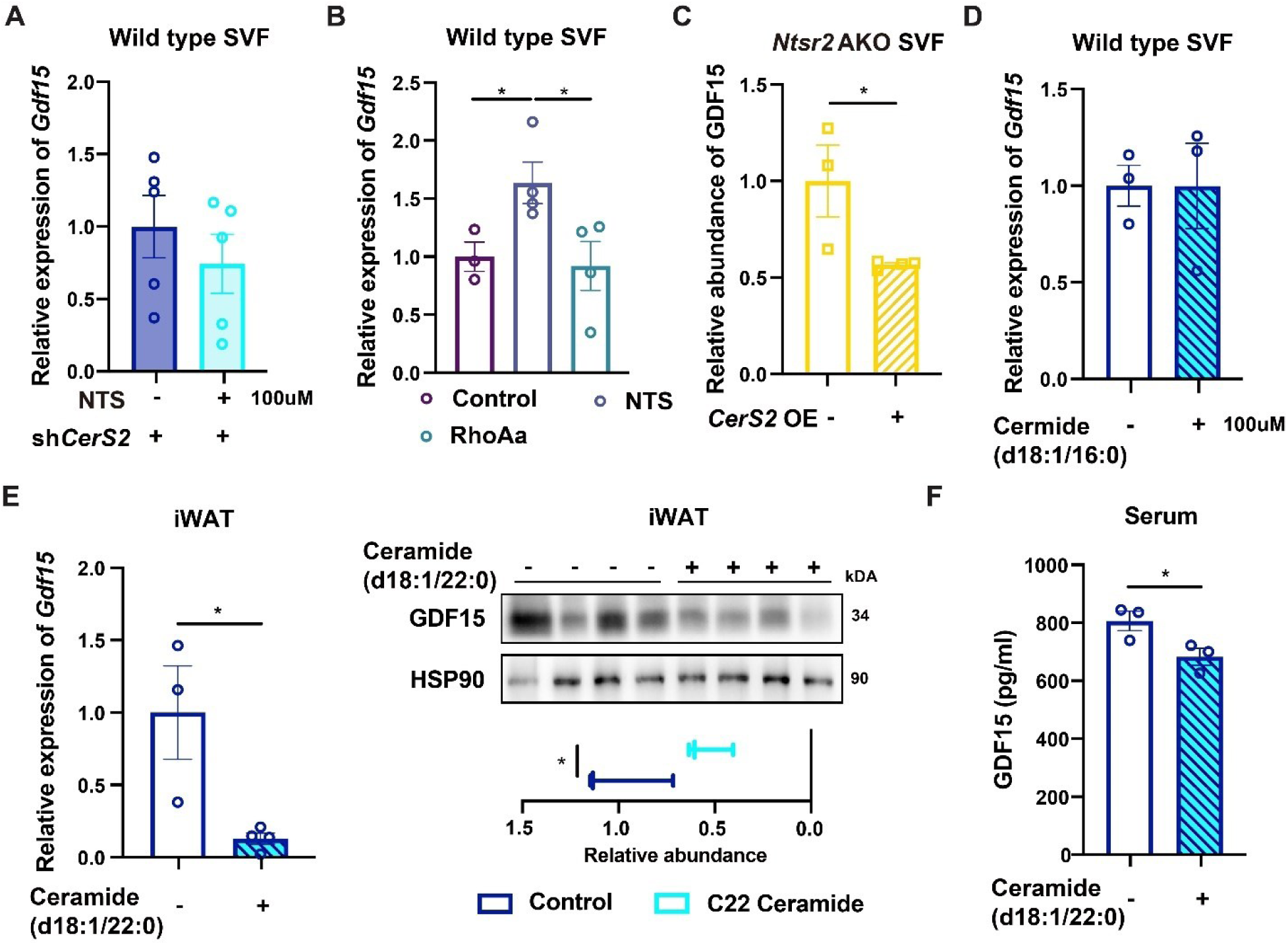
CerS2 is important for NTS-NTSR2 signaling. **A.** The relative expression of *Gdf15* upon NTS treatment in the primary adipocytes with knocking-down of *CerS2*; N=5; **B.** The relative expression of *Gdf15* upon the combinational treatment of NTS and RhoAa in the primary adipocytes. N=3-4; **C.** The production of GDF15 protein in the primary adipocytes with *Ntsr2* AKO and with or without *CerS2* overexpression; N=3-4; **D.** The expression of *Gdf15* gene in the WAT upon the treatment of ceramide C16 *in vivo*. N=3; **E.** The expression of *Gdf15* gene and GDF15 in the WAT upon the treatment of ceramide C22 *in vivo*. N=3-4; **F.** The abundance of GDF15 protein in the serum upon ceramide C22 treatment *in vivo*. N=3. *, p<0.05.

